# Distinct progressions of neuronal activity changes underlie the formation and consolidation of a gustatory associative memory

**DOI:** 10.1101/2021.08.08.455552

**Authors:** Elor Arieli, Nadia Younis, Anan Moran

## Abstract

Acquiring new memories is a multi-stage process. Ample of studies have convincingly demonstrated that initially acquired memories are labile, and only stabilized by later consolidation processes. These multiple phases of memory formation are known to involve modification of both cellular excitability and synaptic connectivity, which in turn change neuronal activity at both the single neuron and ensemble levels. However, the specific mapping between the known phases of memory and the observed changes in neuronal activity remains unknown. Here we address this unknown in the context of conditioned taste aversion learning by continuously tracking gustatory cortex (GC) neuronal taste responses from alert rats in the 24 hours following a taste-malaise pairing. We found that the progression of neuronal activity changes in the GC depend on the neuronal organizational level. The population response changed continuously; these changes, however, were only reflected in the population mean amplitude during the acquisition and consolidation phases, and in the known quickening of the ensemble state dynamics after the time of consolidation. Together our results demonstrate how complex dynamics in different representational level of cortical activity underlie the formation and stabilization of memory within the cortex.

**Significant Statement:** Memories are formed through a multi-phase process; an early initial acquired memory consolidates into a stable form over hours and days. While the underlying phase-specific molecular pathways are fairly known, the neuronal activity changes during these different phases remain elusive. Here we studied this unknown by tracking cortical neuronal activity over 24h as the taste becomes aversive following association with malaise. We found that that the progression of activity changes is organization-level dependent: The population response changed continuously; the population mean amplitude was time-locked to the acquisition and consolidation phases, and the quickening of the known ensemble state dynamics appear only after consolidation. Our results reveal the complex organizational-level neuronal interactions that underlie the progression of memory formation.

## Introduction

Learning is not a discrete event, but rather a process which evolves over time (Bailey and Kandel, 1993; Dudai, 2004; Aceti et al., 2015; Klinzing et al., 2019). More than a century of human and animal research has identified several important phases in the progression of memory formation (Müller and Pilzecker, 1900; Dudai, 2004; Mcleod, 2013; Klinzing et al., 2019; Haubrich et al., 2020), which prominently include: 1) acquisition of an initial, labile memory that is protein-synthesis independent; and 2) consolidation and stabilization of that memory through a process that requires protein synthesis.

These learning dynamics have been mainly studied using the fear conditioning (FC) paradigm in rodents (Schafe et al., 1999; Li et al., 2005; Runyan and Dash, 2005; Johansen et al., 2011; Aceti et al., 2015; Kida, 2019). In this paradigm, the early memory is rapidly acquired in the hippocampus-entorhinal cortex (HIP-EC) network, and consolidation is achieved by transferring the memory into neocortical networks, thus becoming HIP-independent (Squire and Alvarez, 1995; Buzsáki, 1996, 2015; Eichenbaum, 2000; PW Frankland, 2005; Logothetis et al., 2014; Squire et al., 2015; Genzel et al., 2017; Rothschild et al., 2017; Klinzing et al., 2019; Liu and Kuzum, 2019). It is unclear, however, how this stage-wise process, which is ubiquitous across paradigms (Ji and Wilson, 2007; Ortega-Martínez, 2015; Rothschild et al., 2017; Klinzing et al., 2019), plays out in non-hippocampal forms of learning. .

One such hippocampal-independent learning is conditioned taste aversion (CTA) (Garcia et al., 1955; Dibattista, 1988; Dunn and Everitt, 1988; Gallo et al., 1992; Yamamoto et al., 1994; Bures, J., Bermúdez-Rattoni, F., & Yamamoto et al., 1998), a classical conditioning paradigm in which a novel, innately palatable, taste becomes aversive following pairing with malaise. CTA is subserved by a system that includes (but is not limited to) the pons, amygdala and gustatory cortex (GC) (Yamamoto et al., 1994; Welzl et al., 2001; Bermúdez-Rattoni et al., 2004).

While the behavioral taste avoidance reaction can be elicited within minutes of malaise induction (Parker et al., 1984; Spector et al., 1988) and persists for months (Berman et al., 2003), biochemical studies of the GC suggest the existence of both acquisition and consolidation phases occurring within the 24h immediately post learning: acquisition spans the first 3 hours following training, during which protein synthesis inhibition does not disrupt the expression of early taste avoidance reaction, (Rosenblum et al., 1993; Ferreira et al., 2002; Moguel-González et al., 2008); consolidation happens 4-7h post training, and requires protein synthesis (Ferreira et al., 2002; Martinez-Moreno et al., 2011) and brain-derived neurotrophic factor (BDNF) – a molecular mediator for long-term synaptic plasticity (Ma et al., 2011; Xin et al., 2014). These data suggest a distinct transition between the early memory acquisition and the late memory consolidation, both occurring in GC.

The molecular pathways that underlie the formation and consolidation of memories act, at least in part, to change the synaptic strength connecting neurons (Aizenman et al., 2000; Martin et al., 2000; Bi and Poo, 2001; Dudai, 2002; Tye et al., 2008; Tully and Bolshakov, 2010). In turn, these synaptic strength modulations change the firing rates of the postsynaptic neurons (most importantly, the rates in response to taste stimuli)—a change that is believed to constitute the memory itself and drive the learning-induced behavioral change. The presumed “instability” in the neuronal activity during the early acquisition and consolidation phases regularly caused researchers to wait at least 24 hours before testing for learning-related activity changes (Grossman et al., 2008; Moran and Katz, 2014). In the GC, neuronal activity inspected 24 hours after CTA induction indeed showed firing-rate changes (Yamamoto and Fujimoto, 1991; Yasoshima and Yamamoto, 1998; Moran and Katz, 2014). Interestingly, using CTA acquisition and extinction sequentially revealed that GC single neuron response changes poorly predict the behavioral change, but the population dynamic activity did so with high fidelity (Moran and Katz, 2014). Specifically, it was show that GC ensemble state dynamics become quicker following CTA, and slower with extinction (Moran and Katz, 2014). An appealing hypothesis is that CTA consolidation processes organize ongoing single neuronal changes into coherent and optimized ensemble dynamic responses. If that is the case, we will expect to observe the quicker population dynamics only following the consolidation phase. In addition, several other questions remain open regarding the nature of changes during the hours that follow CTA learning: Are the changes confined to some specific times or are they slowly accumulating? What is the relation between changes in the baseline activity and changes in the responses to the stimulus? What is the relation between neuronal activity changes and the biochemically-defined acquisition and consolidation phases?

Here we answer these questions by continuously recording neuronal activity from the GC for 24 hours before and after a CTA training session in alert behaving rats; by periodically delivering taste stimuli throughout this interval, we were able to examine how the changes in the response of single neurons, population representations and ensemble state dynamics evolve across acquisition and consolidation. Our results show that the changes in the activation of these different organizational levels of the GC progress in distinct dynamics. Specifically, single-neuron responses increased during both the acquisition and consolidation phases (but not during the time between them), population responses were continuously changing across the entire time, and the known quickening of the ensemble state dynamics appear only following consolidation. Our results demonstrate the complex interactions between the different organizational levels of the GC whose distinct progression over time underlie the establishment of a CTA memory.

## Methods

### Animals

Male and female Long-Evans adult rats (aged 3-4 months old; ~250g) were raised in groups of 2-4 same sex littermates in a 12H/12H light/dark cycle with the experiments performed in the light portion of the cycle. Following surgery, the rats were single housed in new cages and given wetted food pallets to help with recovery. Their weight was monitored to ensure proper recovery. Unless otherwise specified the rats had *ad libitum* access to chow and water. Rats were handled for 15 minutes a day for 3 days prior to the experiments in order to habituate them to human touch and reduce stress. All methods and experiments carried out in this study comply with the Tel Aviv University Institutional Animal Care and Use Committee guidelines. All efforts were made to minimize animal suffering.

### Surgery and post-surgery care

#### Anesthesia

Rats were temporarily anesthetized with isoflurane (0.5ml/300g) in an induction box, followed by an intraperitoneal injection of a ketamine-xylazine (KX, 100 and 10 mg/kg, respectively, 1ml/250gr). Supplemental intraperitoneal injections (one-third of the induction dose) were administered as needed.

#### Surgery

The anesthetized rat was placed in a stereotaxic frame, its scalp excised, and holes bored in its skull for the insertion of self-tapping ground screws. A craniotomy hole was drilled above the GC (AP = 1.4mm, ML = 5mm, relative to Bregma), and the dura was removed. A self-manufactured electrode with a micro-drive (Piette et al., 2012; Moran and Katz, 2014) was inserted 0.5 mm above the taste cortex (DV = −4.5 from dura). In addition, two intraoral cannulas (IOC, flexible plastic tubing, AM-Systems) were inserted bilaterally through the oral cavity lateral to the second molar tooth (Phillips and Norgren, 1970). The entire structure that included the electrode and IOCs was covered with dental acrylic.

#### Post-operative treatment

Following surgery, the rats were given subcutaneous injections of antibiotics (5mg/kg of Baytril 5%), pain relievers (1mg/kg of Meloxicam 0.5%) and Saline (10ml/kg) to ensure hydration. The head wound margins were treated with antibiotic cream.

### Behavioral procedure

The experimental procedure is illustrated in Fig. 2A.

#### Acclimation (Acc)

After 7 days of recovery from surgery, rats were put into the experimental chamber where they received *ad libitum* water and food for 3 days of acclimation to the new environment.

#### Habituation (Hab)

Following the 3 acclamation days, the rats were connected to the recording system and the IOC was connected to an automatic, nitrogen pressure-based, taste delivery system. The electrodes and the IOC were connected to a self-made dual commutator that provided uninterrupted, tangle-free, taste deliveries and electrophysiological signals. During the next 3 days the water bottle was removed, and the rats were habituated to drink water from a bottle for 20 minutes in the morning, and then to receive bihourly deliveries of 10 drops of water with 15s inter-taste interval through the IOC for a total of ~5ml per 24h. Control rats received saccharin in the bottle and the IOC deliveries instead of water during these days in order to familiarized them with the saccharin taste and inhibit the CTA learning (a latent inhibition effect) (Lubow, 1973). On the 6^th^ day (last day of Hab) we started recording GC neural activity for baseline measurements. There was no need for a control group that receives saline injection instead of LiCl as it was repeatedly shown that GC neuronal responses do not change significantly 24h after CTA (Moran and Katz, 2014).

#### CTA Training

Both the experimental group (Exp) and the taste-familiarized control group (Fam) went through the exact protocol of CTA training. On 9AM of the 7^th^ day (“Training”) all rats were offered a bottle with 0.15% saccharin solution instead of water for 20min. In the hour that followed, the rats received 3 sessions in which saccharin drops were delivered through the IOC to obtain pre-CTA neuronal responses (20 minutes between sessions, 10 drops of 40μl saccharin 0.15% per session). At the end of the last delivery session the rats were subcutaneously injected with a 0.3M LiCl solution (1% body weight) to induce gastrointestinal malaise.

#### Post-injection IOC protocol (24h)

Following CTA induction, the protocol changes to give sessions consisting of 5 drops of saccharin and 10 drops of water distributed in a random order throughout the session (to reduce possible extinction effects). These liquid delivery sessions were given twice during the first hour, then once every hour for 2h and then once every 2 hours until ~4AM. For the next 6 hours the rats had no access to liquids to increase motivation to drink in the following CTA test. The higher initial rate of taste sessions was aimed to probe quicker changes in the short acquisition phase.

#### CTA Test

on 10AM of the 8^th^ day the rats were given a 2-bottle test with water and 0.15% saccharin for 20min. The level of learning was measured by calculating the aversion ratio defined as the consumption of water divided by the total amount of liquids consumed (saccharin and water). 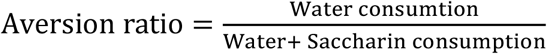

##### Acquisition and analysis of electrophysiological data

Acquisition and pre-processing: Extracellular neuronal signals were collected from self-manufactured 32-wire electrodes (0.0015" formvar-coated Nichrome wire; AM Systems, WA, USA) positioned within the GC (Piette et al., 2012; Moran and Katz, 2014). The data was first collected by an Analog-to-Digital headstage amplifier (RHD2132, Intan Technologies, CA, USA) and then sampled at 30Khz by an Intan RHD2000 acquisition system and stored offline. Common noise was removed from each recorded channel using a common average reference (CAR) algorithm. Automatic spike sorting was first performed using the KlustaKwik python package, followed by manual curation with the Phy program (Rossant et al., 2016). The following stringent criteria were used to ensure that only well isolated and stable neurons will be analyzed: 1) Discriminable action potentials of no less than 3:1 signal-to-noise ratio; 2) Continuous activity across the entire experiment. 3) Clear refractory period of 1ms in the autocorrelation function.

### Perfusion

At the termination of the experiment the rats were anesthetized with KX solution and then perfused with saline (0.9% NaCl) followed by 4% formaldehyde solution. After fixation, the brain was extracted from the skull and left for 72 hours in a 30% sucrose formaldehyde solution at 4℃.

### Histology

The fixed brains were cut to 50μm slices using a microtome (Fisher Scientific), plated on microscope slides, covered with DAPI containing preservative (Invitrogen Flouromount-G with DAPI) and left to dry for 24 hours. Localization of the electrode bundle was accessed using a light binocular. Only rats with correct electrode localization within the GC were included in the study.

### Movement analysis

Full HD (1080p) video recordings were made at 30fps throughout the entire experimental period using Logitech C920 camera. Following experiment termination, the videos were analyzed using the python Open-CV module. The movement of the rat across time was calculated by comparing the differences (in pixels) between adjacent frames of the video. Movement was then averaged over each second (30 frames), and over hours in order to compare between the groups.

### Single neuron response analysis

To study changes in the single GC taste responses across time we divided each neuron’s spiking activity in the 3 secs post a taste event into consecutive 50ms bins. The trials were aggregated into experimental time epochs of pre-CTA, 1H, 2H, 3H, 3-6H, 6-12H and 12-18H post-CTA, and were normalized by dividing the results by the mean BL activity in the 1 second before the aggregated responses. To identify neurons that were changed after 18h we compared pre-CTA and 12-18h taste responses using two-way ANOVA with session (pre-CTA and 12-18h) and response bins (50ms bins between 0-2500ms after taste stimulation) as main factors. Significant (p<0.01) session or interaction effects deemed the neuron as “Changed”, otherwise the neuron was classified as “Unchanged”. Changes in neuronal responses across the experiment time were similarly performed for the early epoch (EE, 200-800ms post taste delivery) and late epoch (LE, 1000-2500ms post taste delivery) response times.

### Assessment of spike train variability at baseline

The Fano factor (FF) is regularly used to assess variability of spike trains. Poissonian Spike trains were first binned into windows of 1s and the spikes in each bin was counted. The FF for every minute of recording during post-CTA time was calculated by dividing the variance of these counts by their mean in non-overlapping windows of 60s.

### Burst analysis

A burst was defined as at least 3 consecutive spikes with inter-spike interval of less than 15ms. Bursts were identified and counted in the 3 seconds before (“BL bursts”) and after (“Response bursts”) each taste event. A burst ratio was calculated by dividing response burst counts by BL bursts counts, that depict the normalized change in bursting activity.

### Ensemble response distance

For each taste delivery event we constructed 3 population FR vectors: baseline, early response and late response. Each was an N-dimensional vector with N being the number of neurons in an ensemble recorded simultaneously from a certain rat. The value assigned to each cell in these vectors was the mean FR during either BL (baseline vector), EE (EE vector) or LE (LE vector). This procedure produced 3 MxN matrices with M the number of taste trials in a certain experimental phase. A reference vector 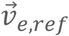 (Fig. 5A), where *e* is one of the epochs (BL, EE or LE), was defined for each matrix by calculating the mean over the 30 Pre-CTA taste events (from the end of saccharin drinking until the LiCl injection). To assess the population changes over time we defined the “Norm distance” 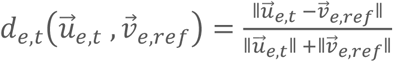; where *e* is one of the epochs, *t* is the time of a block of taste trials, 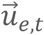 is the mean FR vector of the *e* epoch at block time t, 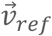 is the mean pre-CTA reference vector and ǁ•ǁ denotes the Euclidean norm of the vector (Fig. 5C). To account for optional drift and BL changes we further normalized the “Norm distance” by performing element-wise subtraction of the BL distance series from the EE and LE series to form the “Normalized Population Response Distance” (NPRD) metric: 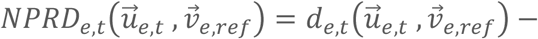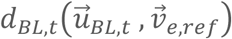, (Notations as defined above). We thus got 3 series of numbers representing the distances from the mean pre-CTA vector (v_ref_) spanning the experimental time.

### Assessing the pattern of LE taste response increases and decreases in the population across time

To understand the relation between LE increases and decreases in the population across time we first calculated the mean and standard deviation of LE responses (1000-2500ms post taste delivery) for each neuron during the Pre-CTA phase. Next, the firing rate in the LE of each trial during post-CTA time was normalized to a z-score by subtracting the mean and dividing by the STD of pre-CTA phase. These values were then averaged for each neuron in each experimental phase. A large negative value indicated a strong decrease compared to pre-CTA, and a large positive value indicated a strong increase. To evaluate the relation between increases and decreases in the population over time we calculated the Pearson correlation between the percentage of neurons with z-score > 1 and those with a z-score < −1.

### Ensemble state dynamics analysis

We assessed the timing of underlying states using a Poissonian hidden Markov models (HMM) (Rabiner, 1989; Kemere et al., 2008; Moran and Katz, 2014). Our model was constructed from 3 consecutive feedforward states; one of the baseline, and two more for the early and late response states. Only a single baseline state was used (instead of 5 like in (Kemere et al., 2008)) since we previously found no difference between the results using different number of baseline states. Trials were grouped into 5 experimental phases: Pre-CTA, 0-3H (CTA acquisition), 3-6H (intermediate), 6-12H (Consolidation) and 12-18H. HMM Training was performed separately for each recorded session and spiking activity of trials from a specific experimental phase. Spike trains were first compiled into 10ms bins to produce momentary spike rate sequences. Probability for maintaining the same state was randomly initialized to values between 0.95 and 0.99 for baseline and early response states, and 1 for the late state. Accordingly, the initial probability of transitioning to the next state in the sequence was 1-P(maintain state) for all states. Firing rates for the baseline, early and late states were initialized to the mean spontaneous rate in the 3 seconds preceding a taste delivery (baseline [BL]), mean firing rate during the first 200-800 ms of post-stimulus time (early epoch [EE]), and mean firing rate during the 1000–2500 ms post-stimulus time (late epoch [LE]), respectively. Training of the HMM was done by application of the forward-backward algorithm; the EM procedure incrementally refined the firing rate and the state transition matrix parameters. We repeated this HMM training procedure 200 times while adding random changes to the initial conditions, eventually using the final state transitions and firing rate matrices with the highest likelihood. We then used the trained model to calculate the posterior probability of the ensemble being in each of the states at each point in time for each trial. At each time bin the ensemble was considered to be in the state which had the highest probability.

### LSTM classifier

A long short-term memory (LSTM) classifier model (Sherstinsky, 2020) was created using Keras and TensorFlow packages in order to reveal the progression of changes in the population (see Fig. 6E). LSTM classifier was used since it uses sequential data as input, and is thus sensitive to the dynamics of the population. Specifically, the LSTM binary classifier was trained on the pre-CTA and 12-18H post-CTA and later was used to classify intermediate phases. The model architecture had an input layer with 32 LSTM nodes which received an input matrix with a shape of 60xN; 60 being the number of 50ms bins in 3 seconds response, and N is the number of neurons. This layer had 15% dropout and 15% recurrent dropout, meaning that 15% of nodes were randomly deleted each training epoch, which was used to facilitate the acquisition of more general features. The LSTM layer was followed by 2 dense hidden layers with 32 and 16 nodes respectively using a ReLU activation function, after which we added Batch Normalization for faster training and a 15% dropout between the layers. The model output layer has 2 nodes with a softmax activation function. The model was trained on a randomly selected set of 70% pre-CTA and 12-18H post-CTA trials, with 15 random model initializations, and tested on the remaining 30% of the trials. The model was optimized using the rmsprop optimizer with ‘binary cross entropy’ and ‘accuracy’ as the loss and metric. Once the model was trained, it was used to predict the probability of each trial from all experimental times being classified as either pre-CTA or 12-18H.

### Statistical analysis

All statistical analyses were done using either a one or two-way ANOVA with a post-hoc analysis using the student’s T-tests. All results are expressed as means + SEM unless stated otherwise. When the results were not normally distributed, a-parametric tests were used; Mann–Whitney for comparing two groups and Kruskal Wallis for comparison between several groups. Data will be available upon reasonable request. Significancy level was set to p<0.05 unless otherwise stated.

## Results

In order to reveal the ongoing progression of GC neuronal activity changes following CTA learning we recorded ensembles of GC single neurons continuously for two consecutive days (48hrs, from 24h before until 24 after CTA training). To achieve this the rats were housed in a custom-built chamber equipped with a combined electric and fluid swivel (Fig 1A), which provided uninterrupted ongoing neural activity monitoring with precisely-timed taste deliveries onto the rat’s oral cavity. Post-experiment electrode position assessment ensured correct localization of the electrode wires within the GC (Fig 1B). The recorded extracellular activity was processed offline to extract and sort neuronal spiking activity. Stringent criteria were employed to ensure that only units clearly isolated from the background noise across the entire recording time will be included in the experimental cohort (Fig 1C, see Materials and Methods section).

**Figure 1:**
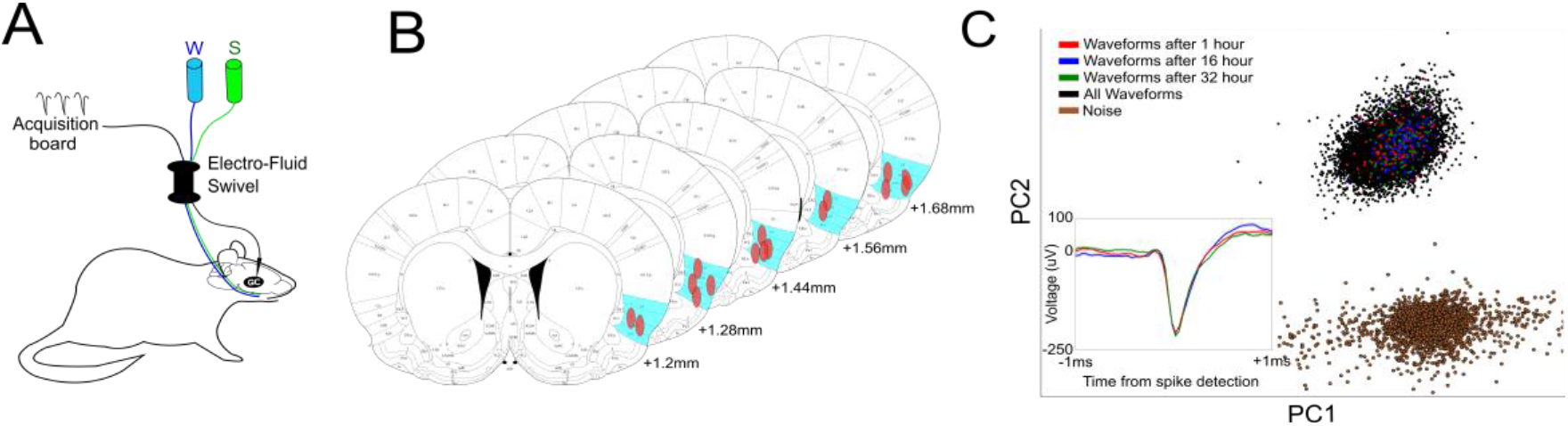
Validation of electrode localization and neuron stability over time. A) Illustration of a rat connected to the experimental apparatus which consists of an Intra-Oral Cannula (IOC) delivering fluids (either water or saccharin) and an SPI cable connected to the implanted electrode in the gustatory cortex on one side and the Intan acquisition board on the other. Both the fluids and electrical cable pass through a dual-swivel component (for the electrical signal and the fluids) in order to allow for free movement of the rat in the chamber. B) localization of the electrode bundle in the gustatory cortex of rats admitted into the experiment marked as a red oval, GC is marked in cyan. C) The shape of all spikes of an example neuron collected across the entire experiment projected on a 2D space following PCA analysis. Different colors indicate waveforms from different timepoints throughout the recording session. Inlet shows the waveform corresponding to each time point. The spike projections are well clustered and far from the noise cluster (brown dot), a good indication of an isolated unit.

### Naïve rats learn CTA but not when familiarized with the conditioned taste

The CTA protocol took 8 days (Fig 2A and the Methods section). Briefly, following acclimation to the recording cage the rats (Exp group, n=7) were habituated to drink water once a day in the morning from two water bottles for 20 minutes, and to receive bi-hourly deliveries of 15 drops of water through the IOC. On the CTA training day (day 7) the rats drunk a novel saccharin solution for 20 minutes from the two bottles, and 60 minutes later received a LiCl injection to induce malaise. Starting immediately after the free saccharin drinking, the rats received scheduled, randomly ordered, intra-oral saccharin and water droplets to monitor neural responses (Fig 2A upper panel, see the Methods section for precise schedule). The level of learning was tested with a two-bottle test in the morning of the 8^th^ day, one containing sacc and the other water. To control for neuronal changes that are not learning-related, we ran an additional group of rats that went through the exact same procedure as the Exp group, but were well-familiarized with saccharin (Fam group, n=7) and therefore should not develop CTA (Lubow, 1973). As expected, both groups drank similar amounts of saccharin during the pre-CTA session on the 7^th^ day (Fig. 2B, 2-sided t-test, t=0.251, p=0.803), but the Exp group showed significantly higher aversion to saccharin than the Fam group after training (Fig 2C, Two-sided t-test, t=4.26, p=0.002). Neural activity across the entire brain, even in primary sensory cotises, is highly affected by motor activity (Musall et al., 2019; Stringer et al., 2019). The use of the Fam control group that received the LiCl also controlled for movement-related biases. Indeed, video-based analysis of the rats’ movement indicated no significant difference between the groups over the post-training hours (Fig. 2D, Two-way ANOVA Group: F(1)=0.16 p=0.68, Group×Time F(1,9)=1.03 p=0.41). Together, these results confirm that while both groups went through the exact CTA procedure, they differ in their CTA learning, thus setting the stage for further examination of the progression of changes in the GC neuronal activity that underlies the learning process.

**Figure 2:**
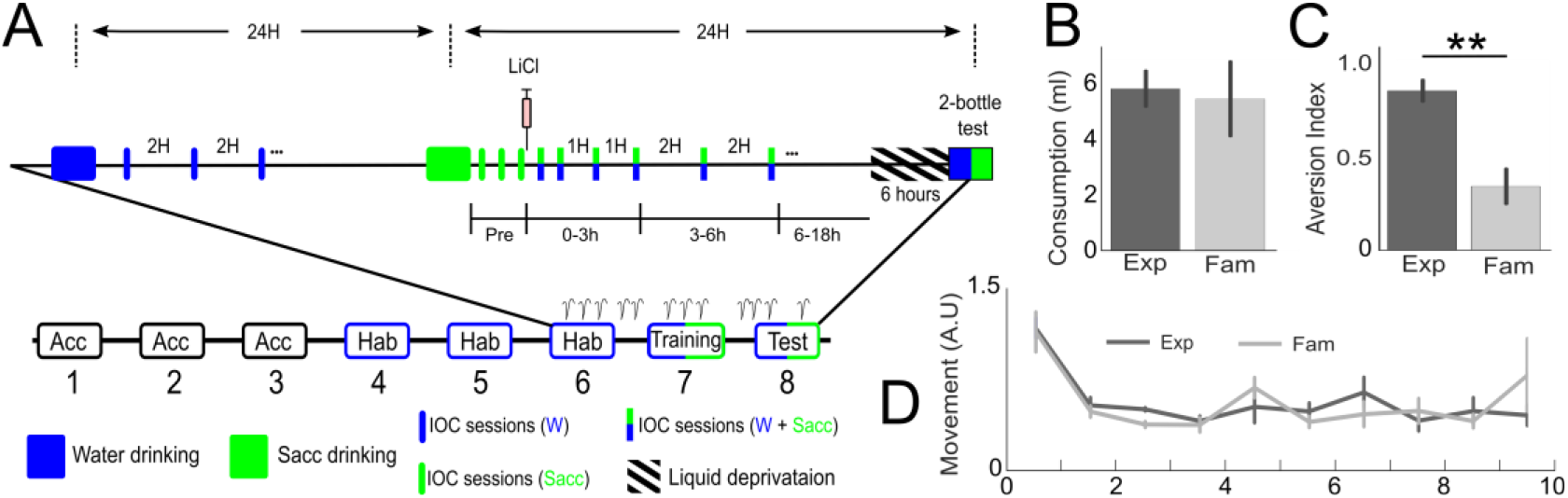
Experimental protocol and behavioral results. A) experimental protocol, rats were implanted with single wire electrode bundles into the gustatory cortex, followed by 1-2 weeks recovery. They were then moved into a custom experimental chamber in which they lived for the duration of the experiment. In the following 3 days the rats acclimated to the new environment. On the 4^th^ day the rats received 10 30ul drops every 2 hours through an IOC to habituate them to the watering protocol. On day 7 (“Training”) the rats received a CTA training by dinking a saccharin solution from the bottles, followed 1h later by a LiCl injection to induce malaise. Following the LiCl injection, the rats received periodic intermixed saccharin and water drops in increasing intervals, starting from 20min and up to 2 hours. Eighteen hours after training IOC deliveries halted for 6 hours. On the 8^th^ day, rats underwent a 2-bottle test to evaluate saccharin aversion. B) The consumption of both Exp and Fam rats was similar on the training day. C) Exp rats show a significantly higher aversion to saccharin on the test day. D) Movement of rats over time, calculated from continuous video recordings of the experimental cage, showed no difference between the groups.

### Learning does not alter the basal neuronal activity properties of the population over time

We recorded 100 and 105 neurons from the Exp and Fam groups (Exp 14.3±3, Fam 15±3.3 neurons per animal), of which 78 and 76 neurons, respectively, were eventually used following stringent excluding criteria of stability and signal to noise ratio. Examples of basal activity across 10 hours from 4 neurons is depicted in figure 3A. These examples show general fluctuation around the mean activity of each neuron, without significant drifts. The mean basal activity of all neurons of Exp and Fam groups showed fluctuation around 8 spikes/second (Fig. 3B), with no significant difference between the groups (Two-way ANOVA, Group F(1)=0.61 p=0.43, Group×Time F(1,9)=1.5 p=0.13). Learning-related changes might not only impact the basal fining rate of neurons, but also the variability of a spike trains (the “regularity” of the spikes in a certain time window). Comparing the Fano factor (FF) that measures that variability, however, revealed no difference between the groups (Fig. 3C, Two-way ANOVA: Group F(1)=0.31, p=0.574, Group×Time F(9)=1.52 p=0.13). Both groups displayed more regular firing rate in the hour following the LiCl injection as indicated by the lower FF values, and a return to a Poissonian pattern (irregular spike sequences, FF values near 1) thereafter (Fig. 3C). The early low FF values are probably related to the LiCl-induced sickness that is known to last for about an hour and is experienced by both groups. Together, the similarity in the movement and neuronal baseline activity characteristics of the two groups provides a good starting point for studying the temporal evolution of learning-related taste response changes.

**Figure 3:**
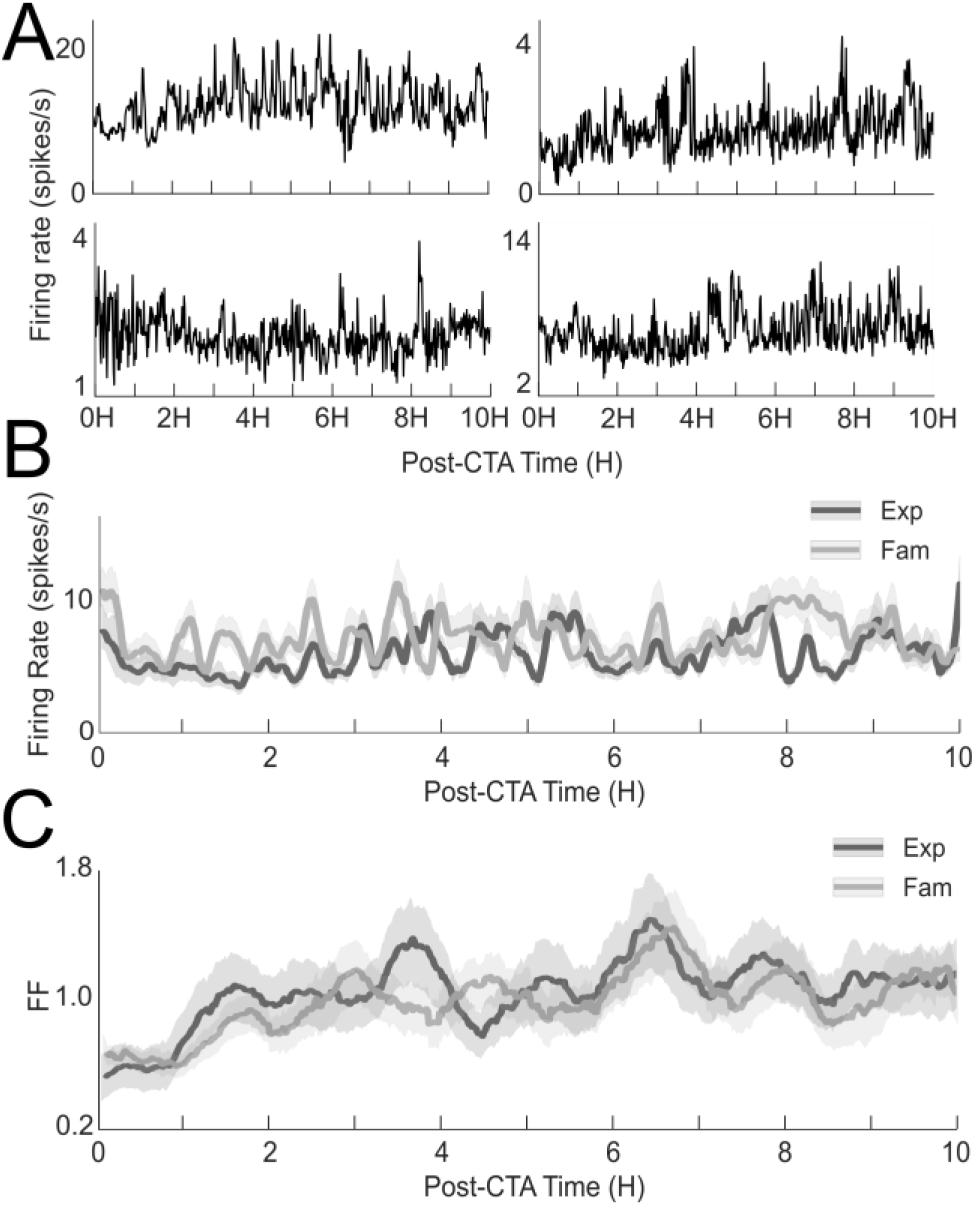
Similar basic neuronal firing rate characteristics between the groups. A) Example firing rates over time with 1min bins of different neurons from both the Exp (left) and Fam (right) rats. B) mean FR of all neurons in both groups over the 10 hours following CTA training. C) Mean Fano-Factor of all neurons in both groups over 10 hours following the LiCl injection. Data are represented as mean ± SEM

### GC neurons change their responses to the conditioned taste following CTA in distinct post-CTA epochs

When tested 18 hours after CTA induction, some of the GC neurons showed altered responses to the saccharin (Fig. 4A upper panel), while others remained unchanged (Fig. 4A lower panel), in accord with previous reports (Moran and Katz, 2014). The response changes were confirmed to be related to the learning; the mean firing rate in response to saccharin during the first 3 seconds increased significantly after 18 hours in the Exp, but not in the Fam group (Fig. 4B left, t-test Exp t(152)=2.85 p=0.006, Fam t(152)=1.345, P=0.181) or the responses to water in both groups (Fig. 4B, right, Two-way ANOVA, Group F(1)=0.97 p=0.32, Time F(1)=0.31 p=0.58 Group×Time F(1,1)=0.87 p=0.35). Recording continuously over the 24 hours following CTA induction allowed scheduled inspection of the progression of neuronal response changes. Based on previous studies we partitioned the 24 hours following CTA induction into several phases: 0-3H (CTA acquisition), 3-6h (intermediate phase), 6-12H(consolidation) and 12-18H (post-consolidation). Figure 3C shows examples of 4 neurons and their progression of response changes over time. While the response of the neuron in Fig. 4C_i_ remained stable over the post-CTA time, the other examples show a diverse pattern of changes, both in the time when changes occurred, and in the direction of changes (increased or decreased). We conducted several more analyses in order to better describe the progression of changes in the single neuron level following CTA induction. Higher rate of taste session in the acquisition phase allowed hourly inspection of changes. Increased percentage of taste responsive neurons in the Exp group was found during the acquisition (2H) and following consolidation phase (6-18H) (Fig. 4D, χ^2^ test: 2H χ(152)=11.64, p<0.001, 6-12H χ(152)=6.48, p=0.011; 12-18H χ(152)=5.58, p=0.018). Additionally, increased neuronal bursting activity known to be associated with learning (Mason and Rose, 1988; Laviolette et al., 2005; Li et al., 2007), was found in the same phases (Fig. 4E, Two-way ANOVA, Group F(1)=16.527 p=0.0006, Time F(6)=3.8 p=0.001 Group×Time F(1,6)=1.66 p=0.13, t-test 3H t(152)=2.82 p=0.006, t-test 6-12H t(152)=2.94 p=0.0048, 12-18H t(152)=4.19, P=0.0001). Another measure that can demonstrate a change in the neuronal response is dissimilarity to the pre-CTA response. We found that the percentages of these neurons increased 2 hours post-CTA, followed by a decline an hour later in both groups (Fig. 4F). However, while these percentages continue to decline in the Fam group during and after the consolidation epoch, they were significantly higher in the Exp group (Fig. 4F, χ^2^ test, 6-12H χ(152)=7.31, p=0.0068); 12-18H: χ(152)=6.34, p=0.012).

**Figure 4:**
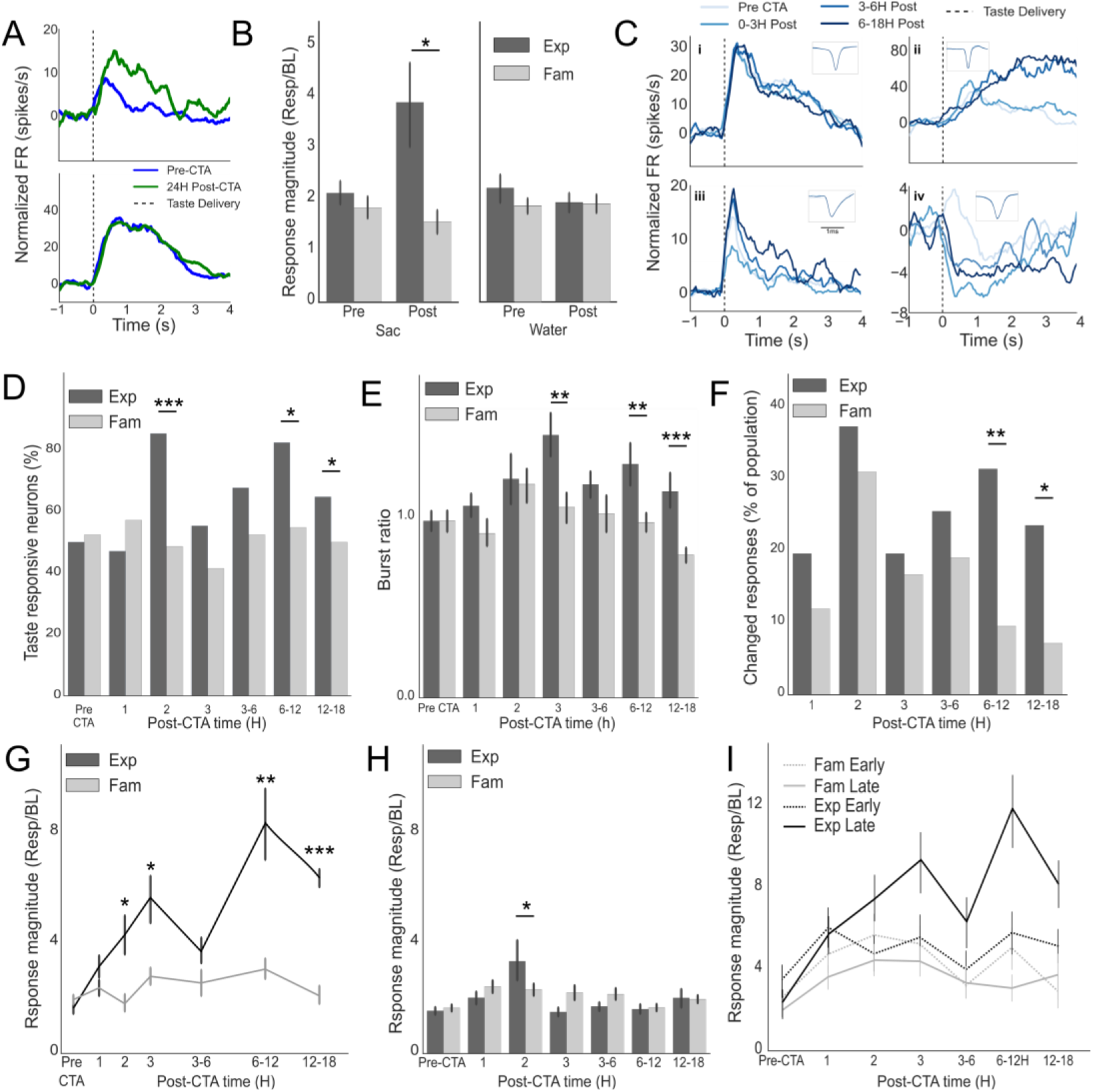
Changes in single neuron responses to the conditioned taste occur in many patterns but are most prominent as a biphasic change. A) Example of a neuron which changed its response to saccharin after CTA learning (upper) and a neuron which didn’t (lower) B). Averaged response magnitude to saccharin (left) increases in GC neurons 18h after CTA in Exp but not in Fam rats. Water response magnitude (right) are unchanged in both groups. C) Example neurons which change their response in various patterns over the hours following CTA training. D) Percentage of taste responsive neurons over the 18 hours post-CTA training. E) Normalized bursting activity during taste responses over post-CTA hours. F) The percentage of neurons which changed their response to the conditioned taste from the Pre-CTA response. Both groups show increased change 2 hours post-CTA, but only neurons of the Exp group have sustained this change 6 hours post-CTA. G) Averaged response amplitude over time of neurons which had significantly different responses 18h post CTA compared to pre-CTA. Neurons in Exp rats show a bi-phasic pattern (2-3 and 6-18 hours post-CTA). H) Population response amplitude over time of neurons which have similar responses between pre-CTA and 18H post CTA. Neurons in Exp rats show a transient increase 2 hours post CTA. I) breakdown of response amplitudes to EE (200-800ms) and LE (1000-2500ms) over hours post CTA shows that changes in the LE amplitude dominated overall amplitude changes over post-CTA time.

How many neurons eventually changed their responses to Sac after consolidation? As expected, we found significantly more such neurons in the Exp group than in the Fam group (Exp: 23.1% [18/78], Fam: 7.9% [6/76], χ^2^ test, χ(152)=6.34, p=0.0012). We considered the neurons that maintained a different response than the pre-CTA after 18 hours as part of the CTA memory engram (Dudai, 2004; Dudai et al., 2015). We wanted to follow the activity of these neurons back to pre-CTA time and reveal the dynamic nature of their changes. To that end we calculated their response magnitude (normalized to baseline, see the Methods section) in different points in time over the post-CTA time. While neurons from the Fam group showed a low and constant response over time, neurons from the Exp group showed double-peak increased responses: one weaker during CTA acquisition and another stronger during and after consolidation (Fig. 4G, Two-way ANOVA, Group F(1)=25.375 p=10^−6^, Group×Time F(1,6)=3.04 p=0.007, t-test: 2H t(22)=2.87 p=0.011, 3H t(22)=2.45 p=0.021, 6-12H t(22)=2.922 p=0.007, 12-18H t(22)=8.52, P=5.26×10^−9^). Interestingly, the neurons that did not change their response following 18h did show a weak but significant response increase 2 hours post-CTA (Fig. 4H Two-way ANOVA, Group F(1)=0.53 p=0.468, Group×Time F(1,6)=2.77 p=0.011, t-test 2H t(128)=2.44 p=0.016). This suggests that neurons that were part of the early, STM-related, part of the learning, may end up out of the final CTA memory engram. Lastly, previous studies showed that taste identity and palatability – the hedonic value of a taste – are coded sequentially in single neuronal responses: an early identity coding epoch (EE, 200-800 msec) is followed by late epoch (LE, 1000-2500 ms) that codes palatability. Since CTA alters the hedonic value of a taste, we expected that the progression of response magnitude changes depicted in Fig. 4G is primarily the result of changes in the LE. When we measured magnitude changes separately for each of the neuronal epochs over time, we found that the double-peak change pattern is the result of LE changes (Fig. 4I), while in the Fam and Exp-EE, the magnitude remains generally unchanged. Together, these results suggest that in the single neuron level of the GC, changes are not accumulating slowly, but rather occur in two distinct epochs: the early acquisition and the consolidation phases; changes which are attributed to alterations in the palatability-coding LE.

### Population response changes start immediately after CTA induction

Inspecting changes in the communal amplitude of single neurons showed a biphasic pattern over the hours following CTA learning, however this type of averaging may hide more delicate changes in the population activation (e.g. balanced excitations and inhibitions that cancel each other out). To study the progression of ensemble-level changes we represented the ensemble activity (separately for the EE and LE) as the sum of N-dimensional vector, with N being the number of neurons recorded simultaneously from each rat (Fig. 5A. Top: EE or LE, bottom: BL, see the Methods section). The BL population activity was similarly represented (Fig. 5A, bottom). Figure 5B shows examples of these vectorized representations over time projected on a 2D plane (using PCA analysis) for Exp (left) and Fam (right) rats. These trajectories, color coded for the experimental time, show the evolution of the population activity changes, both for the LE response (top) and BL activity (bottom). While the Exp response trajectory (top left) appear to deviate more robustly from the initial pre-CTA representation, the trajectory of the Fam example shows unstructured, more random, advancements. To quantify this observation we calculated the “Normalized Population Response Distance” (NPRD, see the Methods section) metric that measures the difference between a population response at a certain phase and the pre-CTA response (Fig 5C, top), normalized to the population BL changes (Fig 5C, bottom). Low values of NPRD (around 0) indicate minor changes in the population. This indeed was the case for the EE (the epoch that codes chemosensory information and not expected to change much) of both Fam and Exp groups, with no significant difference between the two groups (Extended Fig. 5-1, Two-Way ANOVA, Group F(1)=2.7 p=0.09, Group×Time F(1,5)=0.36 p=0.87). In contrast, there was a significant difference between the groups in the palatability-rich LE (Fig. 5D, Two-Way ANOVA, Group F(1)=50.29 p=8.54×10^−22^, Group×Time F(1,5)=9.33, p=4.26×10^−15^). Post-hoc analysis revealed that the LE NPRD was higher in the Exp group compared to the Fam group over the entire post-CTA time (Fig. 5D, 2-sided t-tests; pre p=0.101, 1H p=0.039, 2H p=0.011, 3H p=0.038, 3-6H p=3.3*10^−12^, 6-12H p=0.002). Is this deviation from pre-CTA population response the result of an early single event or the outcome of continuous changes? Investigating the rate of change (the change in the population response compared to the previous session) suggests that the population response kept changing over most of the post-CTA hours in the LE (Fig. 5E, Two-way ANOVA, Group: F(1)=63.1 p=8.06×10^−15^, Group×Time F(1,4)=2.5 p=0.04), but not for the EE (Extended Fig. 5-2). Surprisingly, the population response kept changing during the 3-6H phase (Fig. 5D, E), while the averaged magnitude response of single neurons showed a reduction during this time (Fig. 4H, I). These results suggest that the overall magnitude decline in the intermediate phase might not be a return of single neurons to their pre-CTA responses, but rather the result of some balancing processes. To further study this option, we ordered the neurons in each experimental time according to their Z-score deviation from pre-CTA responses during the LE (Fig. 5F). When we compared the fraction of neurons that changed their responses across the experiment, we found a distinct difference between the two groups: in the Fam group the percentages of neurons that increased and decreased were anti-correlated (r=−0.49, the more increases, the less decreases and *vice versa*), while in the Exp group they were highly correlated (r=0.59, balanced changes of increases and decreases). So far our results suggest that the learning-induced amplitude increases in the LE of GC neurons is the result of continuous changes in the LE ensemble response representation; a progression that is characterized by structured balancing of increases and decreases of the FR in the neuronal population

**Figure 5:**
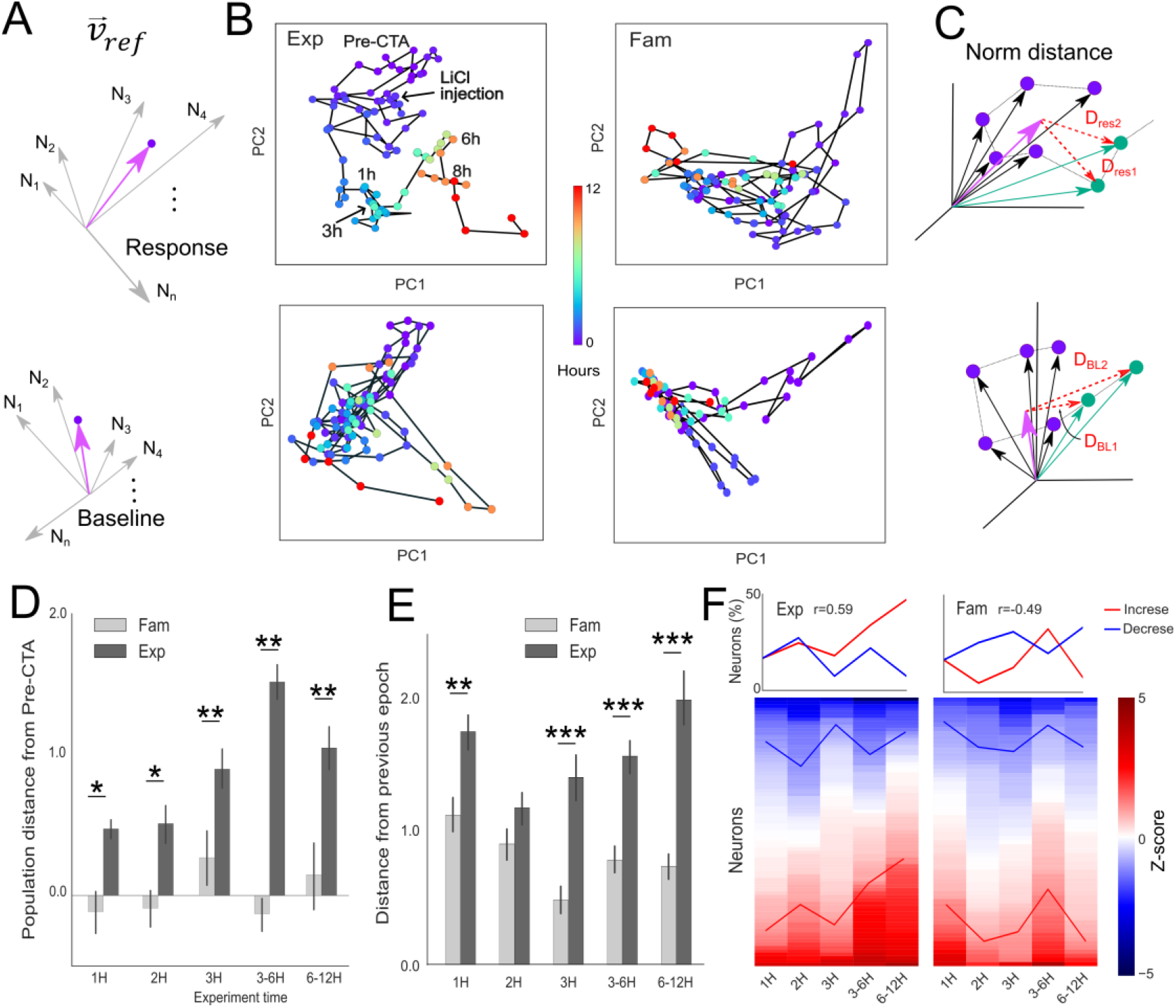
Population responses to the conditioned taste change continuously. A) A vectorized representation of the ensemble activity (top: EE or LE, bottom: BL). Length of arrow indicates FR of a neuron in a certain epoch. Pink arrow indicates the sum of all vectors. Example of ensemble responses to saccharin (top) and BL (bottom) over experimental time, projected on a 2D PCA space, from Exp (left) and Fam (right) rats. Color represents time from CTA induction. While in the Exp rat the ensemble’s response vector seems to increase its distance from the origin, in the Fam rat the changes are more local. C) Illustration of the calculation of the “Norm distance” measure which calculates the Euclidean distance between the vectorize representation of the neuronal firing rate of the ensemble (D_resi_ or D_BLi_) at certain times (green arrows) and the corresponding pre-CTA reference vector V_ref_ (pink arrow). D) NPRD of the LE across time. Exp rats showed increased deviation from pre-CTA values compared to the Fam group across all post-CTA times. E) NPRD calculation as in D, but Vref was set to the preceding phase of each phase tested. This way the calculated values represent the rate of change in the ensemble. The results show that the rate of change in the Exp group is higher than the Fam group during most of the post-CTA experimental time. F) Bottom: A heatmap representation of the response changes in the LE across all neurons in the Exp (left) and Fam (right) groups. Each voxel represents the normalized difference in the LE FR between pre-CTA and a certain time during the experiment, represented in Z-score. Neurons were ordered according to the Z-score values at each time slot. Colored lines represent ±1 z-score values. Top: The Exp group lines shows a correlation between increasing and decreasing responses which indicates a structured change in the network, while neurons from the Fam group show anti-correlation, suggesting a drift effect.

### Learning-induced ensemble state-dynamics quickening occurs only following CTA consolidation

Apart from single neuronal firing rates, taste coding of identity and palatability in the GC were also found in ensemble state dynamics (Jones et al., 2007; Moran and Katz, 2014). Specifically, taste stimuli were shown to elicit sequences of ensemble activity states that were taste specific and faithfully track the hedonic value of the taste such that aversive tastes are processed quicker than palatable tastes. This is also true for learned aversion; measured 24 hours after CTA training, the processing of the conditioned taste becomes quicker (Moran and Katz, 2014). Tracking the progression of ensemble dynamics changes may tell us a great deal about the role they play in CTA learning: if quicker ensemble dynamics appear as early as the single neurons, they will be linked to the aversion behavioral response, while if they appear late (after 6H), they are probably the result of learning-related consolidation processes. To test this we used hidden Markov models (HMM) – a method that was successfully used in previous studies to investigate ensemble dynamics (Abeles et al., 1995; Jones et al., 2007; Kemere et al., 2008; Ponce-Alvarez et al., 2012; Moran and Katz, 2014). For each rat a separate feed-forward 3-state HMM model (Fig. 6A) was trained, one for each experimental phase. Examples of single-trial state probabilities for the same ensemble of neurons across the different experiment times are shown in figure 6B. These examples show that during pre-CTA, 0-3H and 3-6H post-CTA the transition from the EE to the LE occur around 1300-1500 msec after taste experience, but much earlier, around 900 msec, during 6-12H after CTA training (Fig. 6B, bottom). This phenomenon was evident in the Exp group when we compared the averaged state probabilities across trials between pre-CTA (Fig. 6C, left) and 6-12H post-CTA training (Fig. 6C right). When we compare the transition time from the EE to the LE across groups to the experimental time, we found that while transition time remained constant across the experiment in the Fam group (One-way ANOVA, F(3)=1.47, p=0.21), it became significantly quicker than baseline in the Exp group (One-way ANOVA, F(3)=3.4, p=0.017), but only after 6-12 hours post-training (Fig. 6D, 2-sided t-test pre-CTA vs. 6-12h: *t*=2.12, p=0.034). Similar results were found when we used an artificial recurrent neural network to classify ensemble responses as either pre- or 6-12H-post responses. Specifically, we trained a Long short-term memory (LSTM) model classifier using pre- and 12-24h-post ensemble responses to discriminate between the two response patterns (Fig. 6E). The advantage of LSTM over feed-forward models is that it takes into account the temporal structure of the data, and not only its values. After training reached above 95% correct classification, we fed trials recorded at times between these two time points to study the transition patterns. If changes accumulate slowly, we expect to see a steady transition between the start and end times, otherwise a sudden jump is expected. The results of this analysis show these two types of change patterns: while the probability to be classified as “Pre-CTA” tended to decrease steadily with time in the Fam group (Fig. 6 left), it remained high until 6H post training, and then transitioned abruptly to be classified as 12-24H in the 6-12H-post phase (Fig. 6F right), mimicking the HMM analysis results (Fig. 6D). Together, these results suggest that the neuronal FR rate changes that occurred throughout the post-training period (Fig. 4I and 5E), are transformed into changes in the ensemble dynamics only late in the learning process, probably the results of a consolidation-related network reorganization.

**Figure 6:**
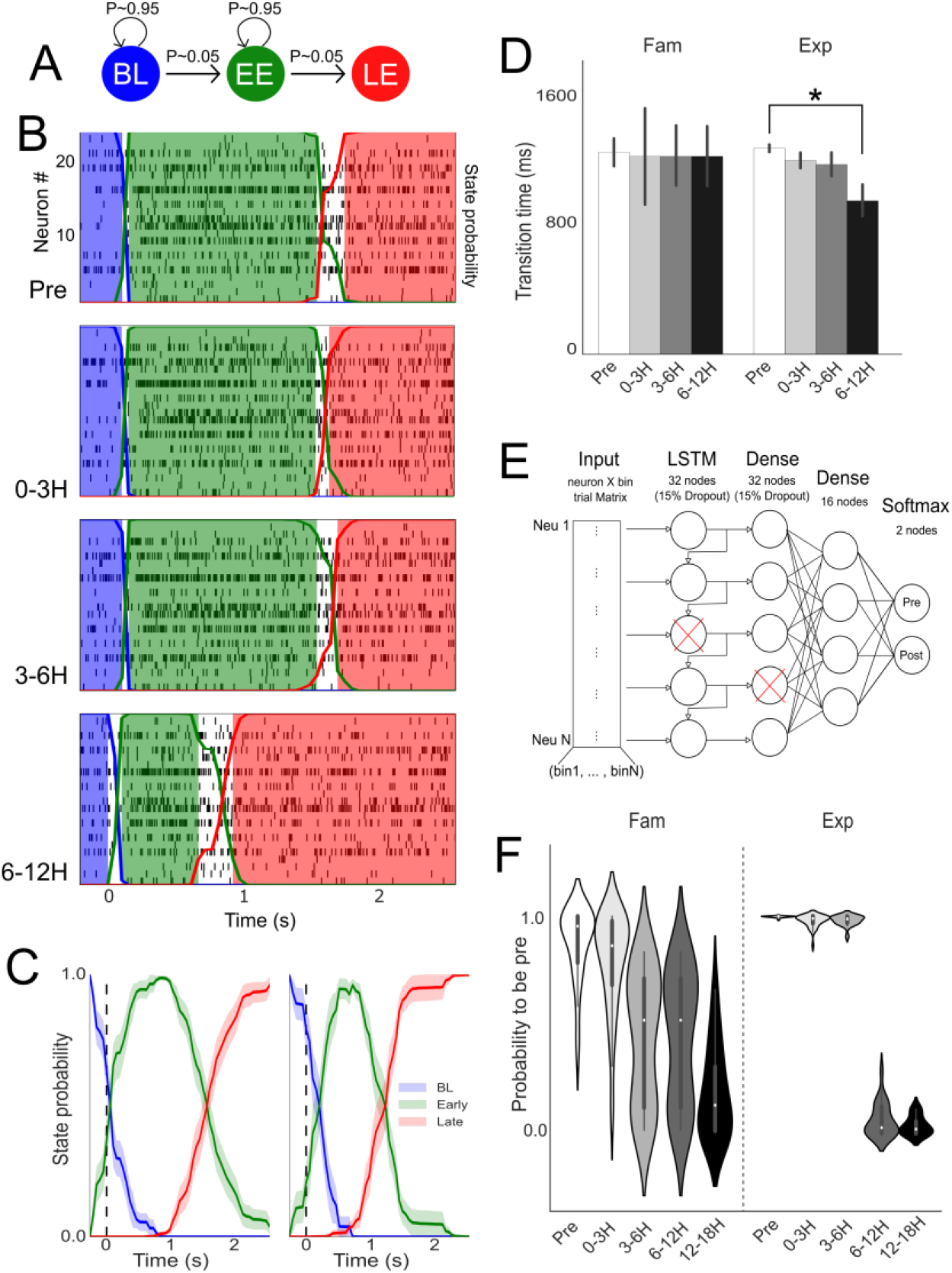
Ensemble dynamics shift abruptly 6 hours post CTA learning. A) illustration of the 3-state transition matrix used for HMM analysis. B) examples over time of state probabilities from an Exp rat. C) mean state probabilities from all Pre CTA trials (left) and all Post CTA trials (right). D) mean transition times from EE to LE states. While in the Exp group we can see a quickening after 6 hours, in Fam rats there was no significant change. E) architecture of the LSTM network used to classify ensemble responses as either Pre or Post CTA. F) Probability to be classified by the LSTM network as either a pre or post CTA trial for trials from different experimental times. While trials from Fam rats show a slow transition between classifications, Exp rats show a clear and abrupt switch 6 hours after CTA learning.

## Discussion

Learning is a dynamic process, subserved by intracellular changes and intercellular connectivity modulations that in turn impact neuronal activity, and consequently behavior behavior (Squire, 1984; Milner et al., 1998; Dudai, 2004). In the taste system, extensive lesion and biochemical research has shown that CTA develops, across the hours following the acquisition trial, from an initially labile memory (Ferreira et al., 2002; Moguel-González et al., 2008) to consolidated learning learning (Ferreira et al., 2002; Ma et al., 2011; Martinez-Moreno et al., 2011). The establishment of a novel memory must necessarily be accompanied by some neuronal activity changes that will generate the appropriate learned reaction; candidate response modulations can be observed early following the taste-malaise association (Yasoshima and Yamamoto, 1998) and 24 hours later (Moran and Katz, 2014), but a systematic understanding of the dynamic progression of learning remains elusive.

Here we directly evaluated neuronal response changes to the conditioned tastant across the first 24 post-training hours, and found that the progression of these changes differs depending on the level of analysis used. At the single neuron level, the mean neuronal response magnitude increased 2-3H post CTA induction, decreased for the next 3 hours (an “intermediate phase”), and increased again 6H post-CTA (Fig 4G): these increased net-activity epochs are a good match for the previously reported acquisition (Ferreira et al., 2002; Moguel-González et al., 2008) and CTA consolidation (Ferreira et al., 2002; Ma et al., 2011; Martinez-Moreno et al., 2011) epochs, respectively; furthermore, the changes were confined to the LE, a palatability-rich part of the GC neuronal responses (Fig. 4I) known to drive reactions to aversive tastes (Mukherjee et al., 2019). Sensitive ensemble-level analyses, meanwhile, revealed different progression patterns across learning. Vectorized representation of the population LE firing rate continued deviating from the pre-CTA representation even in the “intermediate phase” phase (Fig 5E, Extended Fig. 5-2), suggesting that the reduced response amplitude of single neurons during this phase did not reflect reversion to a pre-CTA state. Moreover, we show that despite these ongoing changes in the ensemble neuronal responses, the learning-related quickening of the ensemble state dynamics reported previously 24H following CTA (Moran and Katz, 2014), appear only after the CTA consolidation phase. Together, these three distinct temporal progressions of changes found in the different organizational and functional levels of the GC represent the complex and temporal-specific interactions in which the neurons, and the network they reside within, undergo to support learning.

The evolution of memories and specifically the transition between short- and long-term memory haves been studied for decades (Squire, 1984; McGaugh, 2000; Dudai, 2004). The central role of the HIP in human contextual, semantic, and episodic memory has made the HIP a primary brain region for investigation. Animal studies of these memory systems found an evolution of events that start with an early acquisition in the hippocampus (seconds to hours), an early consolidation phase which changes local hippocampal synapses (synaptic consolidation) and a late (hours to days) consolidation phase (system consolidation) in which the memory is moved to the cortex (e.g., the prefrontal cortex) and becomes more stable, and HIP-independent. This notion, however, does not hold for CTA, in which the HIP is not an essential component (Best and Orr, 1973; Krane et al., 1976; Gallo and Candido, 1995; Yamamoto et al., 1995). Nonetheless, our and previous results suggest that while the dynamics of the learning seems faster in CTA (reaching a consolidated state in less than 24 hours), the memory processes seem to have many similarities, with the early memory acquisition, the involvement of balancing homeostatic processes and the late consolidation. This conserved structure of memory formation, despite differences in its implementation (timing and HIP-dependency) suggests a fundamental property of memory evolution, regardless of the system involved. The difference between the underlying brain circuits of CTA and HIP-dependent learnings might stem from the many “abnormalities” of CTA in the family of classical conditioning learnings. For instance, CTA’s tolerance to long CS-US delays that is supported (at least in part) by GC circuits for taste memory trace formation (Burešová and Bureš, 1980; Welzl et al., 1990; Bermudez-Rattoni, 2014) may involve similar function of the HIP in trace conditioning (Wallenstein et al., 1998; Bangasser et al., 2006). Additionally, the HIP is essential in contextual learning (Kim and Fanselow, 1992; Phillips and LeDoux, 1992; Quinn et al., 2002), a factor which is less important in CTA (Bonardi et al., 1990). Nevertheless, the fact that both CTA acquisition and long-term consolidation happen in the same brain region is intriguing, and calls for further investigation.

It is generally accepted that early memories are labile and then stabilize during consolidation processes. Mechanistically, the distinction between them relates mainly to protein synthesis: the early short-term memory does not depend on protein synthesis but memory consolidation does (Bailey et al., 2004; Dudai, 2004). Based on this criterion we partitioned the time following CTA into early short-term memory acquisition phase (0-3H), an intermediate phase (3-6H) and consolidation phase (6-12H). The early acquisition phase (0-3H) corresponds with the time when the previously consumed novel taste and early CTA memories are formed through a conjunction of acetylcholine (Miranda et al., 2000; Rodríguez-García and Miranda, 2016), dopamine (Guzmán-Ramos et al., 2010), and glutamate (Rosenblum et al., 1997). Later these processes proceed to the activation of the ERK/MAPK pathways involved in the phosphorylation of NMDA receptors and CamKII (Yiannakas and Rosenblum, 2017). LiCl-induced malaise initiates an immediate increase in glutamate in the GC (Miranda et al., 2002; Ferreira et al., 2005), and an off-line increase in dopamine and glutamate about 90 minutes later (Guzmán-Ramos et al., 2010). Importantly, protein synthesis is not required for the early short-term CTA memory (Rosenblum et al., 1993; Moguel-González et al., 2008; Martinez-Moreno et al., 2011). Much less is known about the late CTA consolidation phase in the GC. A requirement for protein synthesis that starts 5H post-CTA mark its earliest time point (Martinez-Moreno et al., 2011). The fact that the protein synthesis was, at least in part required for the synthesis of BDNF, (Ma et al., 2011; Martinez-Moreno et al., 2011) – a known modulator of memories specifically during consolidation (Bramham and Messaoudi, 2005)– further support the relation of this phase to CTA consolidation. To reduce inter-animal variability regarding the transition time into the consolidation phase, we chose 6H post-CTA as the start time of CTA consolidation.

In our study of neuronal activity, both single neurons and ensemble analyses shed light on how brains process and store sensory information, and how learning shapes these memories. In the GC, taste qualities such as taste presence, identity, palatability and familiarity are encoded in a sequence of short epochs of single neurons’ taste responses that span the 2-3 seconds following taste experience (Katz et al., 2001; Bahar et al., 2004; Sadacca et al., 2012). Whereas this epochal activity lives also in GC population ensemble dynamics (Jones et al., 2007), a clear distinction was found between the two levels of analysis in relation to changes following CTA learning: ensemble dynamics show high fidelity with behavior, and single neurons response changes do not (Moran and Katz, 2014). These differences led us to hypothesize that the progression of learning might also play out differently in single neurons and ensemble dynamics. In fact, the time course of post-training response changes in single neurons coincides with the CTA acquisition and consolidation phases (Fig. 4D, 4E, 4F, 4G) This, however, was the result of a non-uniform progression in the level of the neuron; neurons displayed different change pattern of their response across the experiment (Fig. 4C), and only their collective mean response was locked to the acquisition and consolidation phases (Fig.4G). Our results from the acquisition phase are in agreement with previous publication showing increased excitability in single GC neurons shortly after CTA induction (Yasoshima and Yamamoto, 1998). CTA consolidation-specific impact on single neurons activity were not studied before, but post-consolidation increases in neuronal activity in other systems has been reported (Thompson et al., 1996; Aton et al., 2009). The relatedness of the GC amplitude increases to the learning is supported by the fact that they were confined to the LE response time (Fig. 4I), a response epoch of GC neurons know to code palatability information (Katz et al., 2001; Sadacca et al., 2012) and change following CTA (Moran and Katz, 2014). Together, the acquisition and consolidation phases seem to induce robust changes in the activity pattern of single GC neurons.

The two memory phases were separated by the intermediate phase where there was a reduction in many parameters of single neurons activity (Fig. 4D-G), most notably the decrease in the overall response magnitude (Fig. 4G). One explanation to this phenomenon could be the return of the excited neurons closer to pre-CTA response levels. Alternatively, it may be the result of more complicated and continuous changes that balance excitations with and inhibitions across the population. Using vectorized representation of the population activity we found that the GC population responses start changing immediately after CTA learning and continue changing even during the intermediate phase (Fig. 5). This result, which supports the second alternative above, might corresponds to cellular and network-level homeostatic plasticity mechanisms that aim to rebalance the early potentiated activity following the early acquisition phase (Maffei and Fontanini, 2009; Turrigiano, 2011; Keck et al., 2017). Interestingly, it was recently shown that synaptic scaling, a form of homeostatic plasticity, occurs at the same epoch and is essential for the transition from general to specific CTA (Wu et al., 2021). Comparing the relation between the number of neurons that increased or decreased their activity over the post-CTA hours further supported this hypothesis: while in the Fam group increases and decreases were anticorrelated (suggesting a drift effect), in the Exp group they were correlated (Fig. 5F), suggesting a structured and balanced change. While this is an appealing explanation, further direct investigation is required to support a specific role of homeostasis processes during the intermediate phase, and its role in the memory formation.

In the past years it has become increasingly clear that the simultaneous dynamic activity of neuronal ensembles plays a vital role in brain functions. In the taste system, coherent transitions between neuronal ensemble activity states have been found to underlie innate taste-related behaviors (Jones et al., 2007; Miller and Katz, 2010; Jezzini et al., 2013; Mukherjee et al., 2019). By 24H after CTA, these ensemble state dynamics have become faster (Moran and Katz, 2014), allowing for earlier detection and rejection of aversive stimuli. Here we show that this phenomenon does not appear until 6 hours after learning (Fig. 6D, F), following the late consolidation phase. It is important to note that rats display an aversive reaction to the taste-CS as early as 10-20 minutes after LiCl injection (Parker et al., 1984; Spector et al., 1988). Therefore, the faster ensemble dynamics are probably not related to the aversion itself, or its associated motor actions, but rather to a new rearrangement of the network’s connectivity and neuronal excitability. Our data suggests that in the GC, the role of the CTA consolidation phase is to adjust the network’s parameters (neuronal excitability and connectivity) so that the ensemble’s state dynamics become quicker. While previous studies demonstrated the distinction between changes in the single neuron level and the ensemble following learning (Grewe et al., 2017), these were constrained to firing-rate-based coding of the US and CS. Our study, on the other hand, emphasizes the ensemble’s dynamics as the pinnacle of the brain’s target.

The ensemble states sequences found in the taste system (Jones et al., 2007; Moran and Katz, 2014; Mazzucato et al., 2015) other sensory systems (Ponce-Alvarez et al., 2012), the motor system (Abeles et al., 1995; Kemere et al., 2008), and high cognitive brain systems (Balaguer-Ballester et al., 2011) can be naturally modeled as activity transitions in attractor networks (Hopfield, 1982; Amit, 1989). In attractor networks the activity of the nodes quickly shifts between meta-stable states, rather than slowly ramping as information is integrated (Miller and Katz, 2010, 2013; Sadacca et al., 2016). Conceptually, the attractor network can be represented as a landscape with some local dips denoting the network’s states (Fig. 7) (Rolls, 2010). With some external stimulus (such as a taste or a tone), the network activity is pushed to take a certain trajectory within this landscape; but while the inter-trial sequence of states is mostly preserved, the transition times vary and depend on many factors including the network’s activity, history, and noise level. Using HMM techniques, previous studies revealed exactly this “hopping” network dynamics in GC ensemble activity that explained the observed trial-to-trial behavioral variability (Jones et al., 2007; Miller and Katz, 2010; Moran and Katz, 2014; Mazzucato et al., 2015; Sadacca et al., 2016). Importantly, in attractor networks, different network architectures and neuronal activity patterns can produce similar network output (such as reaching one of several “decision” states in the state-space landscape (Fig. 7B and 7C). This fact may explain the ongoing changes in the single-neuron and ensemble activity that still drive the same rejection behavior; while the network’s landscape keeps changing, the CS stimulus will still push the network to reach the “Reject”. The different network configurations may drive a similar reject behavior (Fig. 7B, C) in distinct dynamics. As we know from previous studies, CTA speeds-up ensemble state dynamics for faster ejection of the CS taste (Moran and Katz, 2014). Our current results show that this network phenotype displaying the quickening of network performance, happens at a time known to be related to CTA consolidation (Fig. 6D, F) (Ferreira et al., 2002; Ma et al., 2011; Martinez-Moreno et al., 2011). This result suggests a novel role of consolidation in memory processing: consolidation not only strengthens the memory, making it less labile, but also reshapes the network to change its dynamics (Fig. 7C).

**Figure 7.**
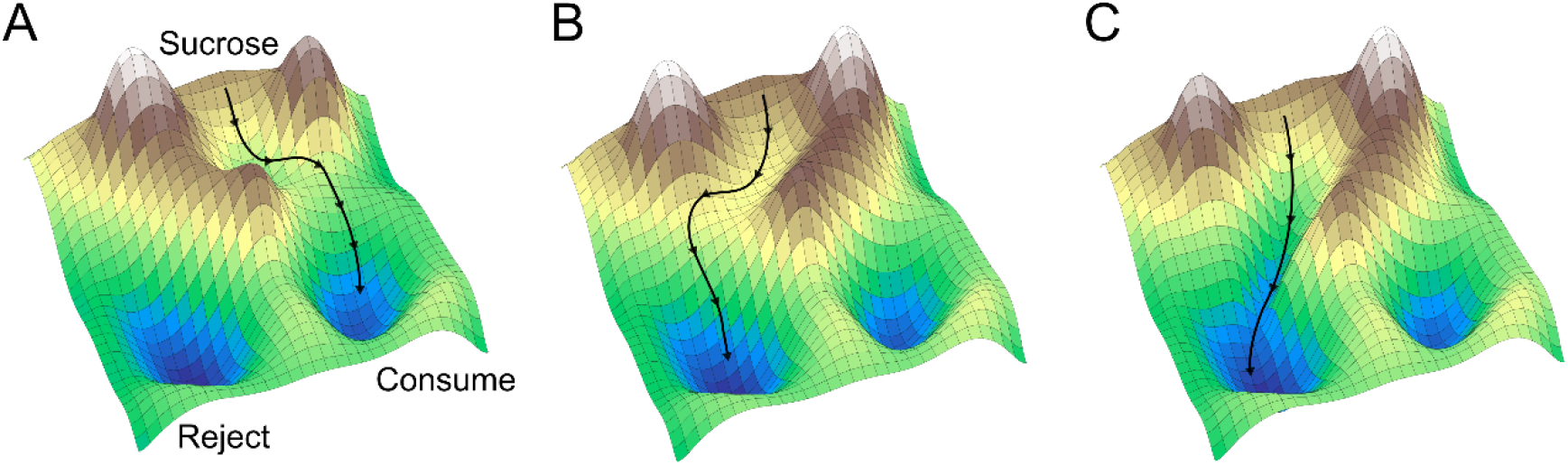
Landscape representation of the neuronal network across learning. The meta-stable states of the network are represented as local dips in the terrain. The neurons’ activity in response to a taste stimulation is some trajectory in the landscape (black line). Prior to learning the network configures such that sucrose stimulation drives the network from the sucrose state to the “consume” decision state. B) Early after training the network configuration changes to divert activity from the sucrose state to the “Reject” state. C) Following consolidation, the network is configured to process sucrose faster in order to quickly reject it upon detection.

Overall, our results highlight the evolution of neuronal changes at different levels of analysis that follow a single trial of taste-malaise associated learning. They highlight the distinct progression of low-level (cellular response amplitudes), mesoscale (ensemble vectorized representations) and higher-level (network dynamics) changes during different phases of learning. These distinct progressions of change are known to underlie complex dynamical system, where the behavior of the system cannot be directly predicted from the behavior of its elements. Nevertheless, further studies are required to elucidate the fine changes in the inter-neuronal connectivity and internal excitability of the neurons that support the creation, stabilization, and maintenance of memories.

## Supplementary material

**Extended Figure 5-1:**
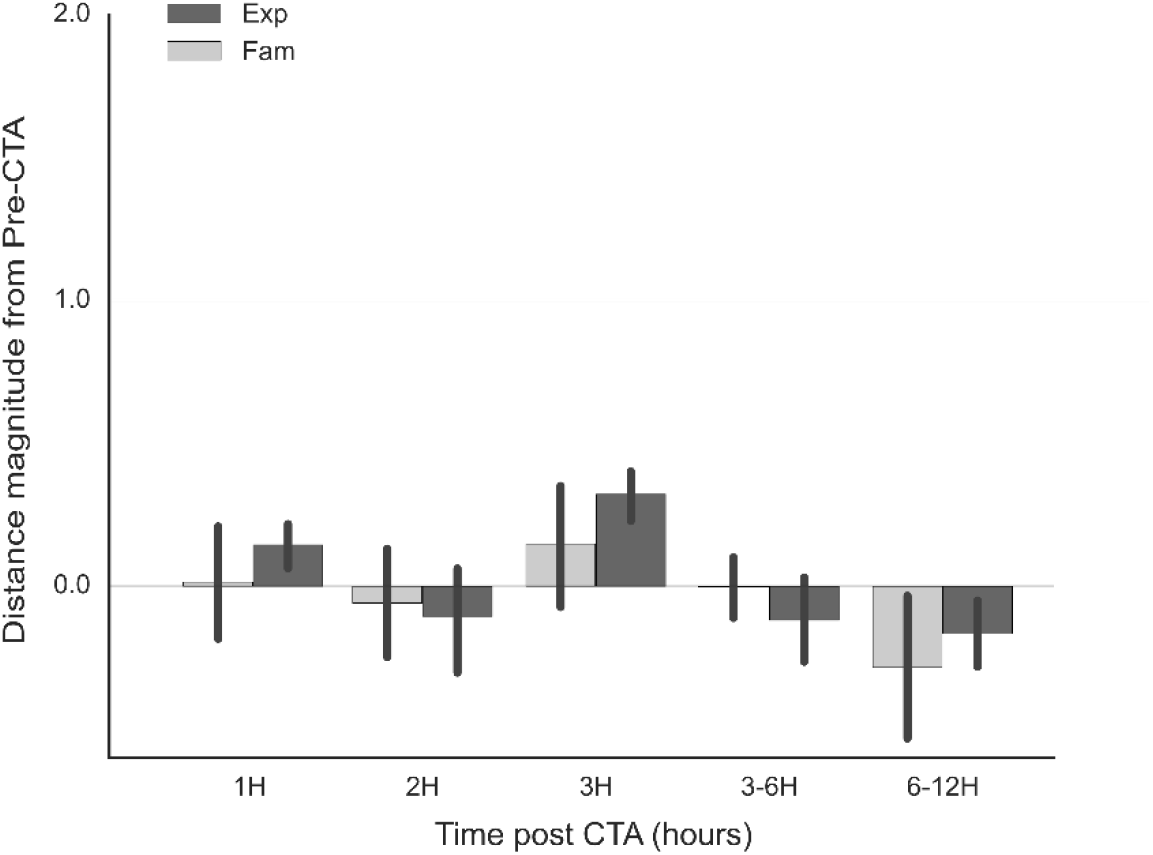
Early epoch population magnitude distance from Pre-CTA across the experiment. The distance magnitude of the population was calculated to evaluate the learning-induced changes the population response, as a whole, across the post-CTA hours. This calculation was performed separately for the early and late response epochs (200-800ms and 1000-2500ms post-stimulus time, respectively. See the methods section). The early epoch of GC neuronal taste responses is known to mainly code chemosensory information, and therefore should not change much following CTA learning. Accordingly, the values of the population distance hovered near 0, indicating overall a stagnant response of the population in this epoch. Also, no significant difference was found between the groups across post-CTA hours (Two-Way ANOVA, Group F(1)=2.7 p=0.09, Group×Time F(1,5)=0.36 p=0.87). Scale of the Y axis was set to match figure 5E in the main article.

**Extended Figure 5-2:**
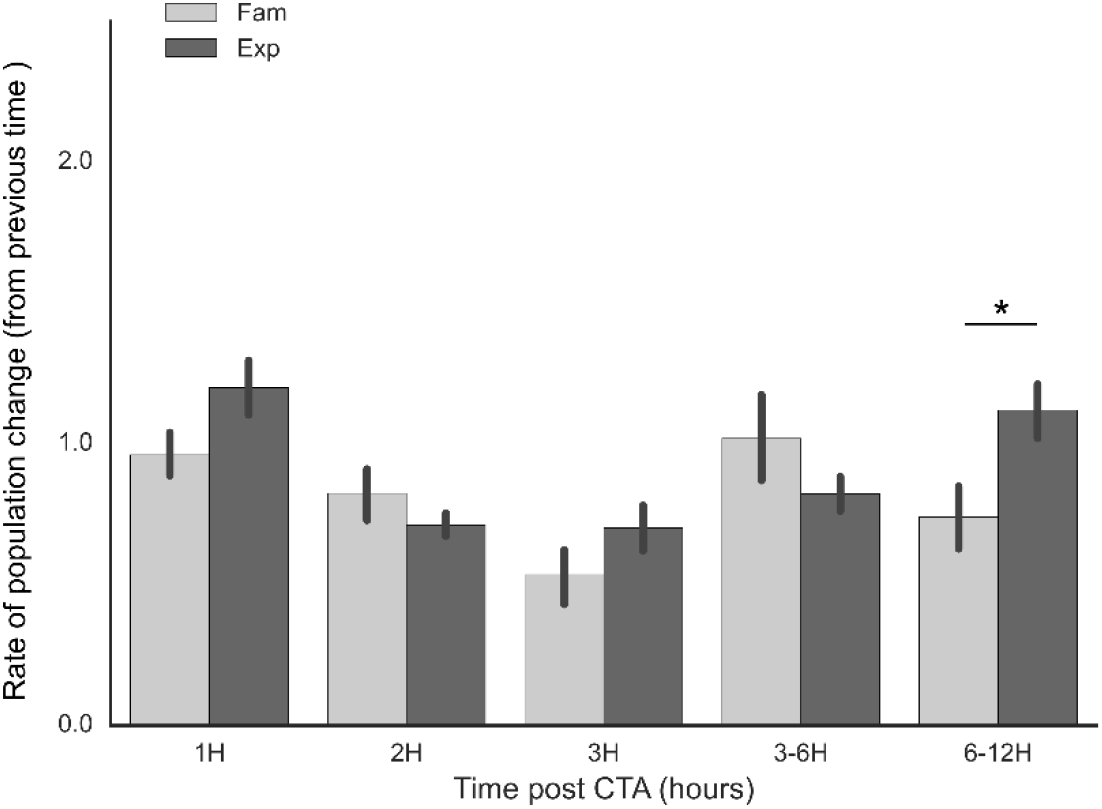
Rate of population response magnitude change over time in the EE. To evaluate the rate of change in the population response we first normalized each population response to its baseline activity, and then calculated the Euclidean distance between the current response and the mean of the previous block (i.e. 1H values were calculated vs pre-CTA, 2H were calculated vs 1H, etc.). Higher values mean a larger change across the population from the previous population response. No significant change between the groups was detected in the EE (Two-way ANOVA, Group: F(1)= 2.98 p=0.08), but there were significant Time and Interaction effects (p=0.0002 and p=0.02, respectively). * p<0.05.

## References

Abeles M, Bergman H, Gat I, Meilijson I, Seidemann E, Tishby N, Vaadia E (1995) Cortical activity flips among quasi-stationary states. Proc Natl Acad Sci U S A 92:8616–20.

Aceti M, Vetere G, Novembre G, Restivo L, Ammassari-Teule M (2015) Progression of activity and structural changes in the anterior cingulate cortex during remote memory formation. Neurobiol Learn Mem 123:67–71.

Aizenman CD, Huang EJ, Manis PB, Linden DJ (2000) Use-dependent changes in synaptic strength at the Purkinje cell to deep nuclear synapse. Prog Brain Res 124:257–273.

Amit DJ (1989) Modeling Brain Function. Cambridge University Press.

Aton SJ, Seibt J, Dumoulin M, Jha SK, Steinmetz N, Coleman T, Naidoo N, Frank MG (2009) Mechanisms of Sleep-Dependent Consolidation of Cortical Plasticity. Neuron 61:454–466.

Bailey CH, Kandel ER (1993) Structural Changes Accompanying Memory Storage. Annu Rev Physiol 55:397–426.

Bailey CH, Kandel ER, Si K (2004) The persistence of long-term memory: A molecular approach to self-sustaining changes in learning-induced synaptic growth. Neuron 44:49–57.

Balaguer-Ballester E, Lapish CC, Seamans JK, Durstewitz D (2011) Attracting Dynamics of Frontal Cortex Ensembles during Memory-Guided Decision-Making Friston KJ, ed. PLoS Comput Biol 7:e1002057.

Bangasser DA, Waxler DE, Santollo J, Shors TJ (2006) Trace conditioning and the hippocampus: The importance of contiguity. J Neurosci 26:8702–8706.

Berman DE, Hazvi S, Stehberg J, Bahar A, Dudai Y (2003) Conflicting processes in the extinction of conditioned taste aversion: behavioral and molecular aspects of latency, apparent stagnation, and spontaneous recovery. Learn Mem 10:16–25.

Bermudez-Rattoni F (2014) The forgotten insular cortex: its role on recognition memory formation. Neurobiol Learn Mem 109:207–216.

Bermúdez-Rattoni F, Ramírez-Lugo L, Gutiérrez R, Miranda MI, Bermudez-Rattoni F, Ramirez-Lugo L, Gutierrez R, Miranda MI (2004) Molecular signals into the insular cortex and amygdala during aversive gustatory memory formation. Cell Mol Neurobiol 24:25–36.

Best PJ, Orr J (1973) Effects of hippocampal lesions on passive avoidance and taste aversion conditioning. Physiol Behav 10:193–196.

Bi GQ, Poo MM (2001) Synaptic modification by correlated activity: Hebb’s postulate revisited. Annu Rev Neurosci 24:139–166.

Bonardi C, Honey RC, Hall G (1990) Context specificity of conditioning in flavor-aversion learning : Extinction and blocking tests. 18:229–237.

Bramham CR, Messaoudi E (2005) BDNF function in adult synaptic plasticity: The synaptic consolidation hypothesis. Prog Neurobiol 76:99–125.

Bures, J., Bermúdez-Rattoni, F., & Yamamoto T, Bures J, Bermúdez-Rattoni F, Yamamoto T (1998) Conditioned taste aversion: Memory of a special kind. (Mackintosh NJ, Shallice T, Schacter D, Treisman A, Weiskrantz L, eds). Oxford: Oxford University Press.

Burešová O, Bureš J (1980) Post-ingestion interference with brain function prevents attenuation of neophobia in rats. Behav Brain Res 1:299–312.

Buzsáki G (1996) The hippocampo-neocortical dialogue. Cereb Cortex 6:81–92.

Buzsáki G (2015) Hippocampal sharp wave-ripple: a cognitive biomarker for episodic memory and planning. Hippocampus 25:1073–1188.

Dibattista D (1988) Conditioned taste aversion produced by 2-deoxy-D-glucose in rats and hamsters. Physiol Behav 44:189–192.

Dudai Y (2002) Molecular bases of long-term memories: a question of persistence. Curr Opin Neurobiol 12:211–216.

Dudai Y (2004) The neurobiology of consolidations, or, how stable is the engram? Annu Rev Psychol 55:51–86.

Dudai Y, Karni A, Born J (2015) The Consolidation and Transformation of Memory. Neuron 88:20–32.

Dunn LT, Everitt BJ (1988) Double dissociations of the effects of amygdala and insular cortex lesions on conditioned taste aversion, passive avoidance, and neophobia in the rat using the excitotoxin ibotenic acid. Behav Neurosci 102:3–23.

Eichenbaum H (2000) A cortical-hippocampal system for declarative memory. Nat Rev Neurosci 1:41–50.

Ferreira G, Gutierrez R, De La Cruz V, Bermudez-Rattoni F (2002) Differential involvement of cortical muscarinic and NMDA receptors in short- and long-term taste aversion memory. Eur J Neurosci 16:1139–1145.

Ferreira G, Miranda MI, De La Cruz V, Rodríguez-Ortiz CJ, Bermúdez-Rattoni F (2005) Basolateral amygdala glutamatergic activation enhances taste aversion through NMDA receptor activation in the insular cortex. Eur J Neurosci 22:2596–2604.

Gallo M, Candido A (1995) Dorsal hippocampal lesions impair blocking but not latent inhibition of taste aversion learning in rats. Behav Neurosci 109:413–425.

Gallo M, Roldan G, Bureš J (1992) Differential involvement of gustatory insular cortex and amygdala in the acquisition and retrieval of conditioned taste aversion in rats. Behav Brain Res 52:91–97.

Garcia J, Kimeldorf DJ, Koelling RA (1955) Conditioned Aversion to Saccharin Resulting from Exposure to Gamma Radiation. Science (80-) 122:157–158.

Genzel L, Rossato JI, Jacobse J, Grieves RM, Spooner PA, Battaglia FP, Fernández G, Morris RGM (2017) The Yin and Yang of Memory Consolidation: Hippocampal and Neocortical. PLoS Biol 15.

Grewe BF, Gründemann J, Kitch LJ, Lecoq JA, Parker JG (2017) Neural ensemble dynamics underlying a long-term associative memory. Nat Publ Gr.

Grossman SA, Fontanini A, Wieskopf JS, Katz DB (2008) Learning-related plasticity of temporal coding in simultaneously-recorded amygdala-cortical ensembles. J Neurosci 28:2864–2873.

Guzmán-Ramos K, Osorio-Gómez D, Moreno-Castilla P, Bermúdez-Rattoni F (2010) Off-line concomitant release of dopamine and glutamate involvement in taste memory consolidation. J Neurochem 114:226–236.

Haubrich J, Bernabo M, Baker AG, Nader K (2020) Impairments to Consolidation, Reconsolidation, and Long-Term Memory Maintenance Lead to Memory Erasure. Annu Rev Neurosci 43:297–314.

Hopfield JJ (1982) Neural networks and physical systems with emergent collective computational abilities. Proc Natl Acad Sci U S A 79:2554–2558.

Jezzini A, Mazzucato L, La Camera G, Fontanini A (2013) Processing of hedonic and chemosensory features of taste in medial prefrontal and insular networks. J Neurosci 33:18966–18978.

Ji D, Wilson MA (2007) Coordinated memory replay in the visual cortex and hippocampus during sleep. Nat Neurosci 10:100–107.

Johansen JP, Cain CK, Ostroff LE, Ledoux JE (2011) Leading Edge Review Molecular Mechanisms of Fear Learning and Memory. Cell 147:509–524.

Jones LM, Fontanini A, Sadacca BF, Miller P, Katz DB (2007) Natural stimuli evoke dynamic sequences of states in sensory cortical ensembles. Proc Natl Acad Sci U S A 104:18772–18777.

Katz DB, Simon SA, Nicolelis MA (2001) Dynamic and multimodal responses of gustatory cortical neurons in awake rats. J Neurosci 21:4478–4489.

Keck T, Hübener M, Bonhoeffer T (2017) Interactions between synaptic homeostatic mechanisms: an attempt to reconcile BCM theory, synaptic scaling, and changing excitation/inhibition balance. Curr Opin Neurobiol 43:87–93.

Kemere C, Santhanam G, Yu BM, Afshar A, Ryu SI, Meng TH, Shenoy K V (2008) Detecting neural-state transitions using hidden Markov models for motor cortical prostheses. J Neurophysiol 100:2441–2452.

Kida S (2019) Reconsolidation/destabilization, extinction and forgetting of fear memory as therapeutic targets for PTSD. Psychopharmacology (Berl) 236:49–57.

Kim JJ, Fanselow MS (1992) Modality-specific retrograde amnesia of fear. Science (80-) 256:675–677.

Klinzing JG, Niethard N, Born J (2019) Mechanisms of systems memory consolidation during sleep. Nat Neurosci 22:1598–1610.

Krane R V., Sinnamon HM, Thomas GJ (1976) Conditioned taste aversions and neophobia in rats with hippocampal lesions. J Comp Physiol Psychol 90:680–693.

Laviolette SR, Lipski WJ, Grace AA (2005) A subpopulation of neurons in the medial prefrontal cortex encodes emotional learning with burst and frequency codes through a dopamine D 4 receptor-dependent basolateral amygdala input. J Neurosci.

Li Y, Zhou W, Li X, Zeng S, Liu M, Luo Q (2007) Characterization of synchronized bursts in cultured hippocampal neuronal networks with learning training on microelectrode arrays. Biosens Bioelectron 22:2976–2982.

Li Z, Zhou Q, Li L, Mao R, Wang M, Peng W, Dong Z, Xu L, Cao J (2005) Effects of unconditioned and conditioned aversive stimuli in an intense fear conditioning paradigm on synaptic plasticity in the hippocampal CA1 area in vivo. Hippocampus 15:815–824.

Liu X, Kuzum D (2019) Hippocampal-Cortical Memory Trace Transfer and Reactivation Through Cell-Specific Stimulus and Spontaneous Background Noise. Front Comput Neurosci 13:67.

Logothetis N, Eschenko O, Murayama Y, Evrard HC (2014) Hippocampal-cortical interaction during periods of subcortical silence.

Lubow RE (1973) Latent inhibition. Psychol Bull 79:398–407.

Ma L, Wang DD, Zhang TY, Yu H, Wang Y, Huang SH, Lee FS, Chen ZY (2011) Region-specific involvement of BDNF secretion and synthesis in conditioned taste aversion memory formation. J Neurosci 31:2079–2090.

Maffei A, Fontanini A (2009) Network homeostasis: a matter of coordination. Curr Opin Neurobiol 19:168–173.

Martin SJ, Grimwood PD, Morris RG (2000) Synaptic plasticity and memory: an evaluation of the hypothesis. Annu Rev Neurosci 23:649–711.

Martinez-Moreno A, Rodriguez-Duran LF, Escobar ML (2011) Late Protein Synthesis-Dependent Phases in CTA Long-Term Memory: BDNF Requirement. Front Behav Neurosci 5:61.

Mason RJ, Rose SPR (1988) Passive avoidance learning produces focal elevation of bursting activity in the chick brain: amnesia abolishes the increase. Behav Neural Biol 49:280–292.

Mazzucato L, Fontanini a., La Camera G (2015) Dynamics of Multistable States during Ongoing and Evoked Cortical Activity. J Neurosci 35:8214–8231.

McGaugh JL (2000) Memory--a century of consolidation. Science (80-) 287:248–251.

Mcleod S (2013) Memory, Encoding Storage and Retrieval. Simply Psychol:1–2.

Miller P, Katz DB (2010) Stochastic transitions between neural states in taste processing and decision-making. J Neurosci 30:2559–2570.

Miller P, Katz DB (2013) Accuracy and response-time distributions for decision-making: linear perfect integrators versus nonlinear attractor-based neural circuits. J Comput Neurosci.

Milner B, Squire LR, Kandel ER (1998) Cognitive neuroscience and the study of memory. Neuron 20:445–468.

Miranda MI, Ferreira G, Ramirez-Lugo L, Bermudez-Rattoni F (2002) Glutamatergic activity in the amygdala signals visceral input during taste memory formation. Proc Natl Acad Sci U S A 99:11417–11422.

Miranda MI, Ramirez-Lugo L, Bermudez-Rattoni F (2000) Cortical cholinergic activity is related to the novelty of the stimulus. Brain Res 882:230–235.

Moguel-González M, Gómez-Palacio-Schjetnan A, Escobar ML (2008) BDNF reverses the CTA memory deficits produced by inhibition of protein synthesis. Neurobiol Learn Mem 90:584–587.

Moran A, Katz DB (2014) Sensory cortical population dynamics uniquely track behavior across learning and extinction. J Neurosci 34:1248–1257.

Mukherjee N, Wachutka J, Katz DB (2019) Impact of precisely-timed inhibition of gustatory cortex on taste behavior depends on single-trial ensemble dynamics. Elife 8:1–32.

Müller GE, Pilzecker A (1900) Experimentelle Beiträge zur Lehre vom Gedächtniss. Z Psychol:1–300.

Musall S, Kaufman MT, Juavinett AL, Gluf S, Churchland AK (2019) Single-trial neural dynamics are dominated by richly varied movements. Nat Neurosci 22:1677–1686.

Ortega-Martínez S (2015) A new perspective on the role of the CREB family of transcription factors in memory consolidation via adult hippocampal neurogenesis. Front Mol Neurosci 8:46.

Parker LA, Hills K, Jensen K (1984) Behavioral eRs elicited by a lithium-or an amphetamine-paired contextual test chamber.

Phillips MI, Norgren RE (1970) A rapid method for permanent implantation of an intraoral fistula in rats. Behav Res Methods Instrum.

Phillips RG, LeDoux JE (1992) Differential Contribution of Amygdala and Hippocampus to Cued and Contextual Fear Conditioning. Behav Neurosci 106:274–285.

Piette CE, Baez-Santiago M a., Reid EE, Katz DB, Moran A (2012) Inactivation of basolateral amygdala specifically eliminates palatability-related information in cortical sensory responses. J Neurosci 32:9981–9991.

Ponce-Alvarez A, Nacher V, Luna R, Riehle A, Romo R (2012) Dynamics of cortical neuronal ensembles transit from decision making to storage for later report. J Neurosci 32:11956–11969.

PW Frankland BB (2005) The organization of recent and remote memories. Nat Rev Neurosci 6:119–130.

Quinn JJ, Oommen SS, Morrison GE, Fanselow MS (2002) Post-training excitotoxic lesions of the dorsal hippocampus attenuate forward trace, backward trace, and delay fear conditioning in a temporally specific manner. Hippocampus 12:495–504.

Rabiner LR (1989) A tutorial on hidden markov models and selected applications in speech recognition. Proc IEEE 77:257–285.

Rodríguez-García G, Miranda MI (2016) Opposing Roles of Cholinergic and GABAergic Activity in the Insular Cortex and Nucleus Basalis Magnocellularis during Novel Recognition and Familiar Taste Memory Retrieval. J Neurosci 36:1879–1889.

Rolls ET (2010) Attractor networks. Wiley Interdiscip Rev Cogn Sci 1:119–134.

Rosenblum K, Berman DE, Hazvi S, Lamprecht R, Dudai Y (1997) NMDA receptor and the tyrosine phosphorylation of its 2B subunit in taste learning in the rat insular cortex. J Neurosci 17:5129–5135.

Rosenblum K, Meiri N, Dudai Y (1993) Taste memory: The role of protein synthesis in gustatory cortex. Behav Neural Biol 59:49–56.

Rossant C, Kadir SN, Goodman DFM, Schulman J, Hunter MLD, Saleem AB, Grosmark A, Belluscio M, Denfield GH, Ecker AS, Tolias AS, Solomon S, Buzski G, Carandini M, Harris KD (2016) Spike sorting for large, dense electrode arrays. Nat Neurosci 19:634–641.

Rothschild G, Eban E, Frank LM (2017) A cortical-hippocampal-cortical loop of information processing during memory consolidation. Nat Neurosci 20:251–259.

Runyan JD, Dash PK (2005) Inhibition of hippocampal protein synthesis following recall disrupts expression of episodic-like memory in trace conditioning. Hippocampus 15:333–339.

Sadacca BF, Mukherjee N, Vladusich T, Li JX, Katz DB, Miller P (2016) The Behavioral Relevance of Cortical Neural Ensemble Responses Emerges Suddenly. J Neurosci 36:655–669.

Sadacca BF, Rothwax JT, Katz DB (2012) Sodium concentration coding gives way to evaluative coding in cortex and amygdala. J Neurosci 32:9999–10011.

Schafe GE, Nadel N V, Sullivan GM, Harris A, LeDoux JE (1999) Memory consolidation for contextual and auditory fear conditioning is dependent on protein synthesis, PKA, and MAP kinase. Learn Mem 6:97–110.

Sherstinsky A (2020) Fundamentals of Recurrent Neural Network (RNN) and Long Short-Term Memory (LSTM) network. Phys D Nonlinear Phenom 404:1–43.

Spector AC, Breslin P, Grill HJ (1988) Taste reactivity as a dependent measure of the rapid formation of conditioned taste aversion: a tool for the neural analysis of taste-visceral associations. Behav Neurosci 102:942–952.

Squire LR (1984) Mechanisms of Memory.

Squire LR, Alvarez P (1995) Retrograde amnesia and memory consolidation: a neurobiological perspective. Curr Opin Neurobiol 5:169–177.

Squire LR, Genzel L, Wixted JT, Morris RG (2015) Memory consolidation. Cold Spring Harb Perspect Biol 7.

Stringer C, Pachitariu M, Steinmetz N, Reddy CB, Carandini M, Harris KD (2019) Spontaneous behaviors drive multidimensional, brainwide activity. Science (80-) 364.

Thompson LT, Moyer JR, Disterhoft JF (1996) Transient changes in excitability of rabbit CA3 neurons with a time course appropriate to support memory consolidation. J Neurophysiol 76:1836–1849.

Tully K, Bolshakov VY (2010) Emotional enhancement of memory: how norepinephrine enables synaptic plasticity. Mol Brain 3:15.

Turrigiano G (2011) Too many cooks? Intrinsic and synaptic homeostatic mechanisms in cortical circuit refinement. Annu Rev Neurosci 34:89–103.

Tye KM, Stuber GD, de Ridder B, Bonci A, Janak PH (2008) Rapid strengthening of thalamo-amygdala synapses mediates cue-reward learning. Nature 453:1253–1257.

Wallenstein G V., Eichenbaum H, Hasselmo ME (1998) The hippocampus as an associator of discontiguous events. Trends Neurosci 21:317–323.

Welzl H, Alessandri B, Bättig K (1990) The formation of a new gustatory memory trace in rats is prevented by the noncompetitive NMDA antagonist ketamine. Psychobiology 18:43–47.

Welzl H, D’Adamo P, Lipp HP (2001) Conditioned taste aversion as a learning and memory paradigm. Behav Brain Res 125:205–213.

Wu C-H, Ramos R, Katz DB, Turrigiano GG (2021) Homeostatic synaptic scaling establishes the specificity of an associative memory. Curr Biol:1–12.

Xin J, Ma L, Zhang T-Y, Yu H, Wang Y, Kong L, Chen Z-Y (2014) Involvement of BDNF Signaling Transmission from Basolateral Amygdala to Infralimbic Prefrontal Cortex in Conditioned Taste Aversion Extinction. J Neurosci 34:7302–7313.

Yamamoto T, Fujimoto Y (1991) Brain mechanisms of taste aversion learning in the rat. Brain Res Bull 27:403–406.

Yamamoto T, Fujimoto Y, Shimura T, Sakai N (1995) Conditioned taste aversion in rats with excitotoxic brain lesions. Neurosci Res 22:31–49.

Yamamoto T, Shimura T, Sako N, Yasoshima Y, Sakai N (1994) Neural substrates for conditioned taste aversion in the rat. Behav Brain Res 65:123–137.

Yasoshima Y, Yamamoto T (1998) Short-term and long-term excitability changes of the insular cortical neurons after the acquisition of taste aversion learning in behaving rats. Neuroscience 84:1–5.

Yiannakas A, Rosenblum K (2017) The insula and taste learning. Front Mol Neurosci 10:1–24.

